# Benchmarking sketching methods on spatial transcriptomics data

**DOI:** 10.1101/2025.08.26.672376

**Authors:** Ian K. Gingerich, Brittany A. Goods, H. Robert Frost

## Abstract

High-throughput spatial transcriptomics (ST) now profiles hundreds of thousands of cells or locations per section, creating computational bottlenecks for routine analysis. Sketching, or intelligent sub-sampling, addresses scale by selecting small, representative subsets. While effective for scRNA-seq data, existing sketching methods, which optimize coverage in expression space but ignore physical location, can introduce spatial bias when applied to ST data. To explore the impact of sketching on ST analysis, we systematically benchmarked uniform sampling, leverage-score sampling, Geosketch (minimax/Hausdorff), and scSampler (maximin) across multiple real ST datasets (mouse ovary, MERFISH brain, human breast cancer, lung) and simulations, using three input representations: PCA embeddings, spatial coordinates, and spatially smoothed embeddings. We show that expression-only designs capture global transcriptomic heterogeneity but distort tissue architecture by over-sampling high-variability regions and under-sampling homogeneous areas. Coordinate-only sampling restores tissue coverage but misses transcriptional extremes. A simple spatially aware extension, computing leverage scores from a randomized SVD basis smoothed by a spatial weights matrix, strikes a favorable balance, recovering rare cell states while maintaining uniform tissue coverage and avoiding edge effects. Across robust Hausdorff distances, clustering stability (ARI), PCA loading drift, and local cell-type MSE, spatially smoothed leverage scores match or outperform alternatives. These results motivate joint spatial–transcriptomic sketching objectives to enable fast, unbiased analyses of increasingly large ST datasets.

## 1 Introduction

### 1.1 Sketching

As high-throughput genomic assays have driven datasets into the millions of observations and thousands of features, intelligent down-sampling, or sketching, has become a core strategy for scalable analysis, visualization, and method development. Rather than sampling uniformly at random, principled sketching techniques aim to preserve the geometric and statistical structure of the data to maintain coverage of rare states, boundary regions, and transitional trajectories [1–3]. Design-based (explicit sampling-design) approaches formalize this goal: minimax criteria (often expressed via Hausdorff distance) choose subsets that minimize the worst-case distance from any point to the sketch, controlling coverage gaps, whereas maximin criteria maximize the minimum pairwise distance among selected points, promoting well-spread and non-redundant samples [4]. In addition to influence-based (importance-weighted or leverage/score–driven) schemes such as leverage score sampling, these strategies provide general-purpose recipes for constructing small, representative sketches that support exploration, inference, and benchmarking across large-scale datasets [5].

### 1.2 Sketching for scRNA-seq data

Single cell RNA-seq (scRNA-seq) datasets are increasingly analyzed with “sketching” or intelligent sub-sampling: selecting a representative subset of cells that preserves transcriptomic diversity while dramatically reducing computational load. A naive approach is uniform random sampling, which is simple and unbiased with respect to sampling probability but often fails to capture rare cell types and transitional states, especially when data are highly imbalanced. Uniform sampling also tends to over-represent dense regions in the gene-expression manifold and under-represent sparsely populated regions, leading to loss of important biological variation. Additionally, it has been demonstrated that over-sampled cell types tend to be over clustered, ie: a single biological population is artificially split into multiple small, highly similar clusters driven by local density fluctuations, technical noise, or batch effects, while under-sampled cells are under clustered, ie: distinct rare or transitional populations are merged into a single broad cluster because there are too few points to resolve clear boundaries [6]. Down-sampling can also improve the signal-to-noise ratio in a dataset by reducing random correlations caused by variation in data quality and read-mapping bias [7, 8]. To address these issues, existing non-uniform methods seek to sample evenly across transcriptomic space: they identify or construct a low-dimensional representation (e.g., PCA, latent embeddings), quantify each cell’s “coverage” or influence within that space, and then preferentially sample points that increase coverage of underrepresented regions while de-emphasizing redundant points in dense clusters. In scRNA-seq, such approaches, including leverage score-based sampling, minimax/Hausdorff-based geometric sketching, and maximin designs, have become standard tools for creating compact yet faithful summaries of large datasets [5, 9–15].

### 1.3 Sketching for ST data

High-resolution array-based platforms such as 10x Genomics Visium HD produce measurements at millions of subcellular-sized locations per slide; in practice, data are commonly aggregated into roughly 300, 000–600, 000 spatial bins to balance resolution and computational tractability, with each bin carrying transcriptome-wide counts for 20, 000+ genes. In situ hybridization–based platforms such as 10x Genomics Xenium routinely profile on the order of 200, 000–500, 000 cells per tissue section, with larger studies exceeding half a million cells, across targeted panels of 5, 000 genes. As these technologies mature, multi-section studies and atlases now comprise hundreds-of-thousands to millions of spatial locations, each with high-dimensional gene expression and precise spatial coordinates [16–26]. While the scale of these datasets promises unprecedented views into tissue architecture, it also creates a big-data bottleneck: standard exploratory workflows, clustering, visualization, neighborhood enrichment, spatial domain detection, become prohibitively slow and memory-intensive without principled sub-sampling strategies that respect both transcriptomic and spatial structure.

Despite their successful use for scRNA-seq analysis, current sketching algorithms operate solely on the gene expression manifold and ignore physical location. One widely used strategy, implemented in Seurat [5], computes approximate statistical leverage scores for each cell using an efficient randomized algorithm. These leverage scores are then converted into sampling probabilities for sketching. This approach is motivated by the fact that cells with high leverage are those that exert strong influence on the embedding and higher sampling probabilities for these cells enriches for transcriptomic diversity [5, 11]. Geometric sketching methods further formalize the objective of achieving approximately uniform coverage of the data manifold by controlling point-wise approximation error and reducing coverage gaps across regions of varying density. Geosketch [10] and Hopper [12] cast sketch selection as a minimax (Hausdorff) design problem: choose a subset *S* of size *k* that minimizes the maximum distance from any point in the dataset to its nearest neighbor in *S*. Concretely, letting *X* be all points (cells/locations) in an embedding and *S* ∈ *X* the sketch, the (directed) Hausdorff distance from *X* to *S* is the worst-case nearest-neighbor distance, which measures the largest “hole” in coverage. Minimizing this objective ensures that every region in transcriptomic space is close to at least one selected point, limiting coverage gaps that would otherwise exclude rare or transitional cell states. Geosketch uses approximate farthest-point sampling over a grid to efficiently reduce this maximum distance, while Hopper accelerates the search and pruning to further tighten the Hausdorff criterion. A related approach, scSampler [9], adopts a maximin design: it selects points to maximize the minimum pairwise distance within the sketch, spreading selected samples as uniformly as possible across the high-dimensional expression space. Collectively, these methods target the same goal; uniform coverage of transcriptomic variability, via different optimization strategies and implementations. However, transcriptome-only sketching can distort spatial structure in ST data. If high-variability or high-leverage profiles cluster in a particular region of a tissue, a purely expression-based sketch will over-sample that region and under-sample others. In ST data, where neighborhood relationships, cell–cell contact patterns, and spatial gradients of expression are essential for interpretation, such imbalances can systematically bias downstream analysis. Over-representing a spatial hotspot inflates the apparent frequency of certain cell–cell adjacency and co-localization relationships, leading to biased estimates of neighborhood enrichment or depletion and misleading microenvironment niches (Fig. 1A). Methods that integrate ligand–receptor expression with spatial proximity will overestimate interactions prevalent in over-sampled regions and underestimate interactions in under-sampled regions, skewing inferred communication networks and hub cell types. Over-coverage of one region and under-coverage of others distorts estimated spatial trajectories and continuous expression gradients (e.g., along anatomical axes), flattening or exaggerating trends and misplacing boundaries between niches (Fig 1B). Biased sketches also alter the apparent spatial distribution of gene expression and domain boundaries, potentially merging distinct regions or fragmenting coherent tissue compartments due to uneven sampling density (Fig.1C). Because most downstream tasks rely on the same sampled subset, either directly or for building reference embeddings early spatial bias propagates through the entire pipeline, compounding errors in subsequent inferences and reducing reproducibility across datasets and laboratories.

**Figure 1.**
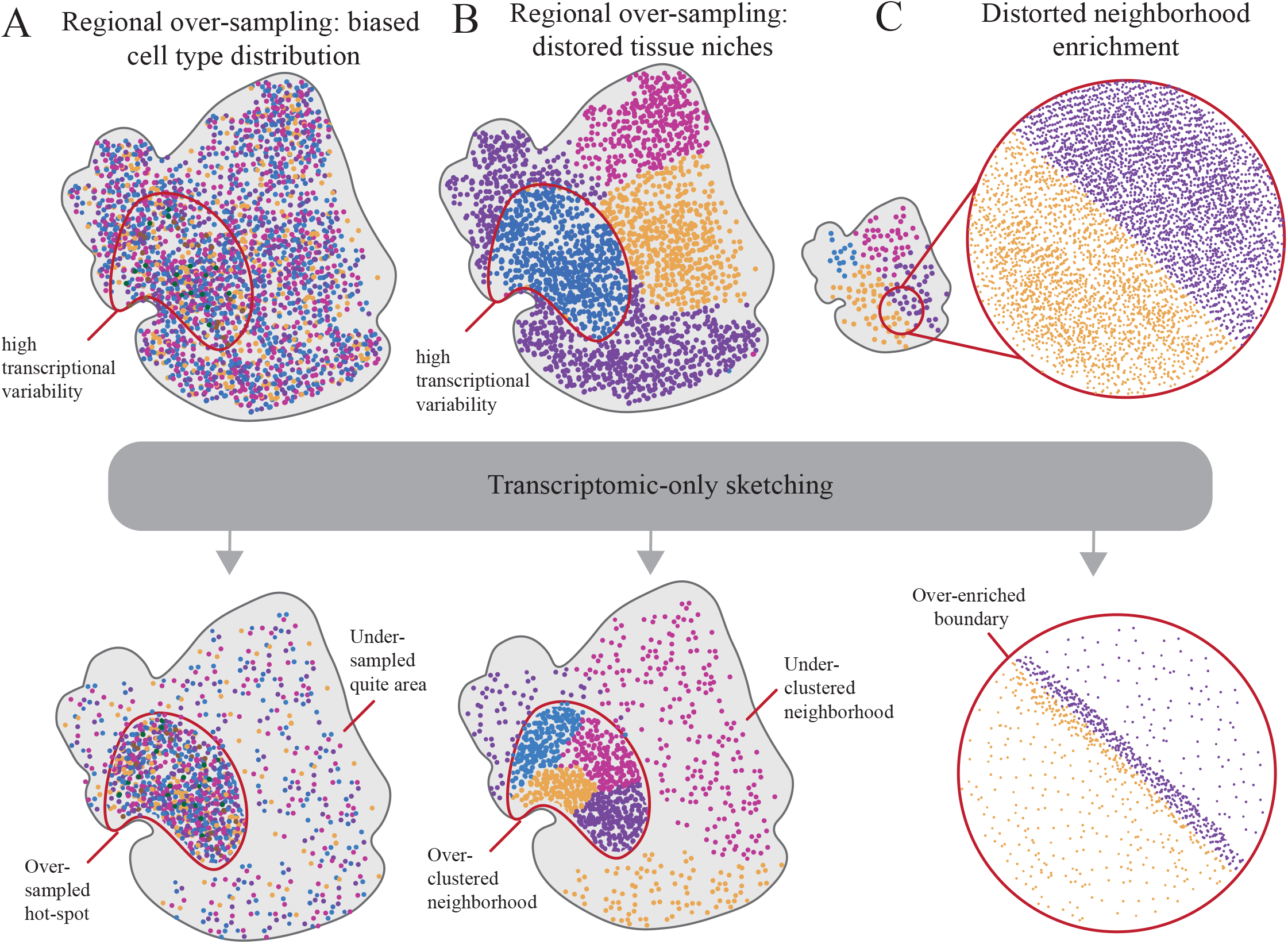
Examples of transcriptomic-based sketching bias. **A:** Regional over-sampling leads to biased cell type proportions and over-representation in transcriptional hotspot. **B:** Regional over-sampling leads to over clustered tissue niches in transcriptional hotspot. **C:** Over-sampling at cell type or neighborhood boarder leads to exaggerated neighborhood enrichment.

To further explore these challenges, we benchmarked current sketching approaches on real and simulated ST datasets to evaluate their ability to preserve both transcriptomic heterogeneity and spatial organization. Our results highlight scenarios in which transcriptome-driven sketches distort tissue architecture and underscore the need for spatially informed sampling strategies. By systematically comparing existing methods on real spatial datasets, we lay the groundwork for adapting sketching algorithms to the unique challenges of ST data, coupling coverage in expression space with balanced coverage in physical space to maintain fidelity for downstream spatial analyses.

## 2 Methods

### 2.1 Evaluated ST datasets

We benchmarked four families of sketching methods (see Table 2) across five public ST datasets spanning multiple platforms and spatial resolutions (MERFISH, Xenium, Visium HD) and six different simulation models. For real data, we performed quality control by excluding cells/locations with fewer than 100 detected counts and removing outliers exceeding five times the mean absolute deviation. After filtering, data were library-size scaled and log-normalized, followed by truncated principal component analysis to 20 components for downstream sketching and evaluation. Real ST datasets included: a mouse ovary MERFISH dataset with eight ground-truth cell-type labels from the original study [27]; a human breast cancer 10x Xenium dataset with histology-derived annotations [28]; a human lung 10x xenium dataset without ground-truth labels, for which we generated full-data reference clusters; a sagittal mouse brain MERFISH dataset [26] with region, class, and type annotations, where ARI was computed against cell-class labels; and a coronal mouse brain 10x Visium HD dataset spanning cortex, subcortex, and hippocampus, evaluated against full-data graph-based clusters due to the lack of ground truth. Simulated data comprised six datasets (three Visium-like on a grid, three Xenium-like with randomly distributed coordinates) with complex, radial, and striped spatial layout (see Table 1 for details). Simulated ST data was generated with SRTsim [29] using a fixed seed and included six label classes (A–F) with specified high/low-signal and noise gene counts, dispersion, baseline mean expression, zero-inflation, and distinct class-specific log-fold changes to induce separable spatial domains (see Table **??**). See Extended Data Section 1.1 for dataset-specific preprocessing details.

**Table 1:**
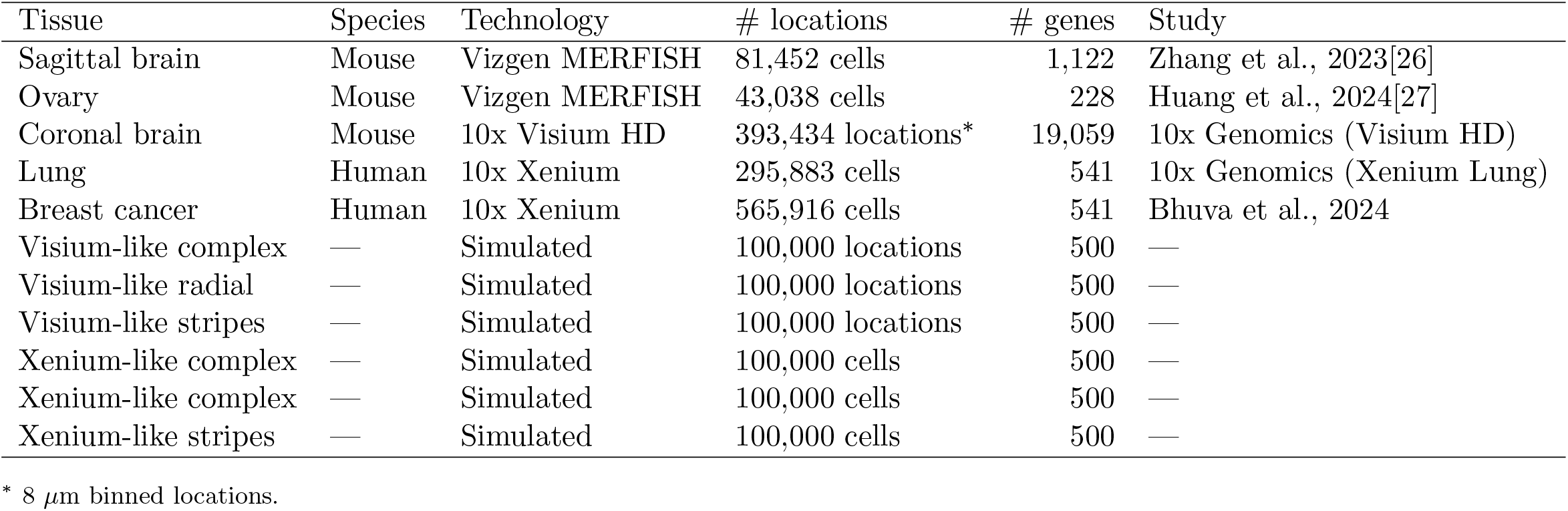
Dataset summary

**Table 2:** Sketching algorithms evaluated and their configurations.

| Algorithm | Input representation | Design/criterion | Language | Study |
| --- | --- | --- | --- | --- |
| scSampler | PCA matrix | Maximin | Python | Song et al., 2022[9] |
|  | Spatial coordinates |  |  |  |
|  | Spatially smoothed PCA |  |  |  |
| Geosketch | PCA matrix | Minimax | Python | Hie et al., 2019[10] |
|  | Spatial coordinates |  |  |  |
|  | Spatially smoothed PCA |  |  |  |
| Leverage score | Cell-by-gene matrix | Statistical influence | R | Clarkson & Woodruff, 2017[5] |
|  | PCA matrix |  |  |  |
|  | Spatially smoothed U |  |  |  |
| Uniform | Cell index | Random sampling | Python/R | — |

### 2.2 Evaluated sketching algorithms

We compared four sketching algorithms: uniform sampling, Leverage score [5], Geosketch [10], and scSampler [9]. Each was applied to three input representations when applicable: (i) transcriptomic embeddings derived from PCA (20 components), (ii) raw spatial coordinates, and (iii) spatially smoothed embeddings that incorporate local neighborhood structure. Uniform sampling selected indices without replacement using equal probabilities and distinct random seeds per replicate. Leverage score methods prioritize statistically influential observations either from the original expression space (variable feature selection as appropriate) or from a low-rank embedding with sampling probabilities were proportional to normalized leverage values. For the standard leverage-score computation, we follow Seurat’s implementation, which uses random projection followed by QR decomposition to approximate leverage scores efficiently on large matrices. For the spatially smoothed leverage-score variant (our simple extension), we first compute a low-rank randomized SVD (rSVD) of the cell-by-gene matrix to obtain *U* (left singular vectors). We then apply spatial smoothing by multiplying *U* by a precomputed spatial weights matrix *W*, yielding *Ū* = *WU*. Leverage scores are computed from the smoothed basis *Ū* (row-wise squared norms), normalized to probabilities, and sampled without replacement. Geometric sketching implements a minimax (Hausdorff coverage) objective to distribute samples so that all regions of the embedding are close to at least one selected point. Maximin designs aim to maximize the minimum pairwise distance among selected points, yielding a uniformly spread subset in high-dimensional space. Spatially smoothed variants use a precomputed spatial weights matrix to smooth the low-dimensional representation prior to sketch selection, thereby encouraging balanced sampling across physical space while preserving major transcriptomic variation. All sketches were drawn without replacement at pre-specified sampling fractions; procedures with inherent randomness were repeated ten times with different seeds to assess stability. See Table 2 for all listed algorithms. See Extend data section 1.2 for details on sketching algorithm design and implementation.

### 2.3 Evaluation metrics

We evaluated each sketch along complementary transcriptomic and spatial criteria. Robust (partial) Hausdorff distance quantified worst-case nearest-neighbor discrepancies between the sketch and full data, computed both in expression space (e.g., PCA) and in physical coordinate space. We used a quantile-based, outlier-robust formulation to reduce sensitivity to extreme points and employed approximate neighbor search to scale to hundreds of thousands of locations. The Adjusted Rand Index (ARI) assessed concordance between sketch-derived clusters and reference labels: ground-truth annotations when available (MERFISH ovary, MERFISH brain class, Xenium breast cancer) or full-data graph-based clusters otherwise (Xenium lung, Visium HD brain). To evaluate preservation of global variance structure, we computed a PCA projection difference by comparing the sketch’s projections onto principal components learned from the full dataset versus components learned from the sketch; lower values indicate more consistent low-dimensional structure. To measure local compositional fidelity, we computed the mean squared error between k-nearest-neighbor cell-type proportion vectors derived from the full versus sketched datasets at each sketched location, summarizing changes in local neighborhood makeup. Finally, we derived an overall spatial score by rank-summing ARI, coordinate-space Hausdorff distance, and local neighborhood distortion at a fixed sampling fraction, emphasizing joint preservation of group structure, spatial coverage, and microenvironmental context. See Extended data section 1.3 for detailed metric computations.

For each sketched dataset, we computed a variety of evaluation metrics to quantify the effectiveness of the associated sketching algorithm. Previous methods have used the transcriptomic Hausdorff distance as their primary or sole method for evaluating sketch quality, however, we feel that this metric is insufficient to characterize the spatial characteristics of the sketched data. To this end, we employed additional metrics to better assess sketch quality[9, 10]. These metrics include the robust Hausdorff distance, Adjusted Rand Index (ARI) between the sketch-based clustering solution and ground truth annotations, mean difference between PCA loading vectors, and distortion of the local neighborhood structure (see Fig 2B and 2C). This evaluation framework provides a framework to understand the impact of different down-sampling strategies on the analysis of ST data, facilitating informed decisions in experimental design and data interpretation. See Section 2.3 for details on evaluation metric calculations. In brief:

**Figure 2.**
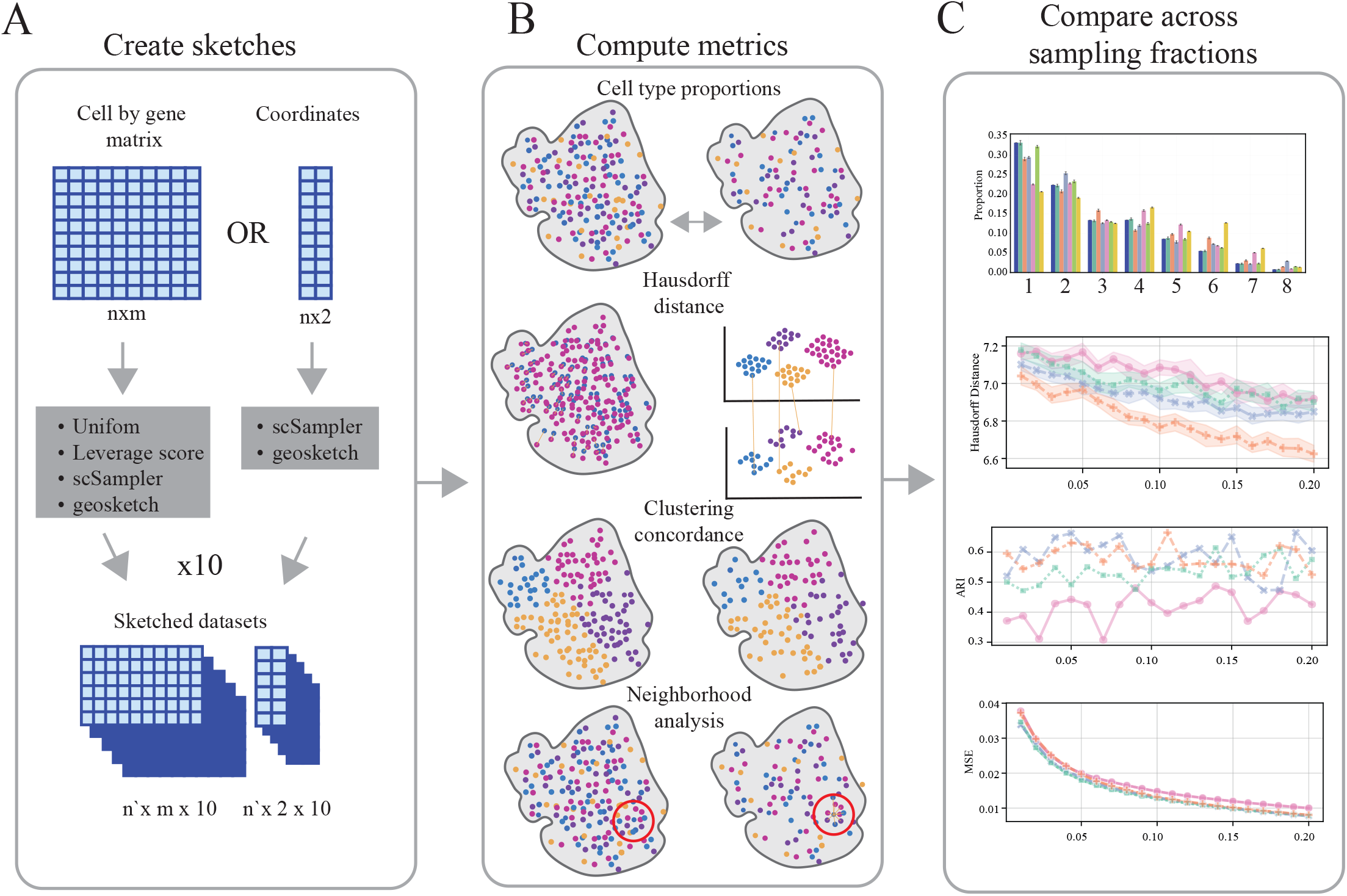
Analysis pipeline. **A:** Sketching protocol **B:** Sketched dataset evaluation metrics **C:** Metric quantification across sampling fractions

- **Robust/partial Hausdorff distance (expression and spatial)**. Quantifies the worst-case nearest-neighbor discrepancy between the sketch and the full dataset, computed separately in transcriptomic space and in physical space. Lower values indicate better coverage (fewer “holes”) and greater fidelity of the sketch to the full data, with the partial version reducing sensitivity to outliers.
- **Adjusted Rand Index (ARI)**. Measures agreement between cluster labels obtained from the sketched data and ground-truth (or full-data-derived) labels, adjusted for chance. Scores range from −1 to 1, with higher values indicating better recovery of biological group structure.
- **PCA projection difference**. Compares how the sketched data project onto PCs learned from the full dataset versus PCs learned from the sketch itself. Smaller mean distances imply that the sketch preserves the dominant variance structure of the full data and yields consistent low-dimensional representations.
- **Local neighborhood distortion (MSE of cell-type proportions)**. Assesses how well the sketch preserves local tissue composition by comparing, for each cell, the kNN-based cell-type proportion vector between the full and sketched datasets. Lower MSE indicates better retention of neighborhood context and microenvironmental structure.
- **Overall spatial score (rank-sum across metrics)**. Aggregates ranks across ARI, coordinate-space Hausdorff, and local neighborhood distortion to summarize spatial fidelity at a fixed sampling fraction. Lower (better) rank-sum reflects methods that jointly preserve group structure, spatial coverage, and local composition.

## 3 Results

Across five diverse tissues, sub-sampling purely on coordinates ensures uniform spatial coverage but pays a price in transcriptomic diversity; sketching purely on gene expression values ensures that transcriptomic extremes are included but biases sampling toward biologically variable sub-regions. These findings motivate the development of sketching methods that explicitly weight the tradeoff between spatial uniformity and transcriptomic diversity.

### 3.1 Sketching result summary

Figs 3 and 4 visualize the overall performance for sketching fractions of 0.1 across all simulated and real ST datasets. For the simulated Visium-HD-like and coronal mouse brain datasets, uniform sampling, leverage score computed on the full cell-by-gene matrix, and leverage score on the spatially smoothed left singular vectors perform well across all evaluation metrics and show a robust overall spatial score (3B (left side of heatmap, columns 1-3) and4D (column 3)). For the other real and simulated datasets, coordinate-based sketching and spatially-smoothed leverage scores result in the best overall score, with the traditional scSampler and Geosketch design preforming the worst.

**Figure 3.**
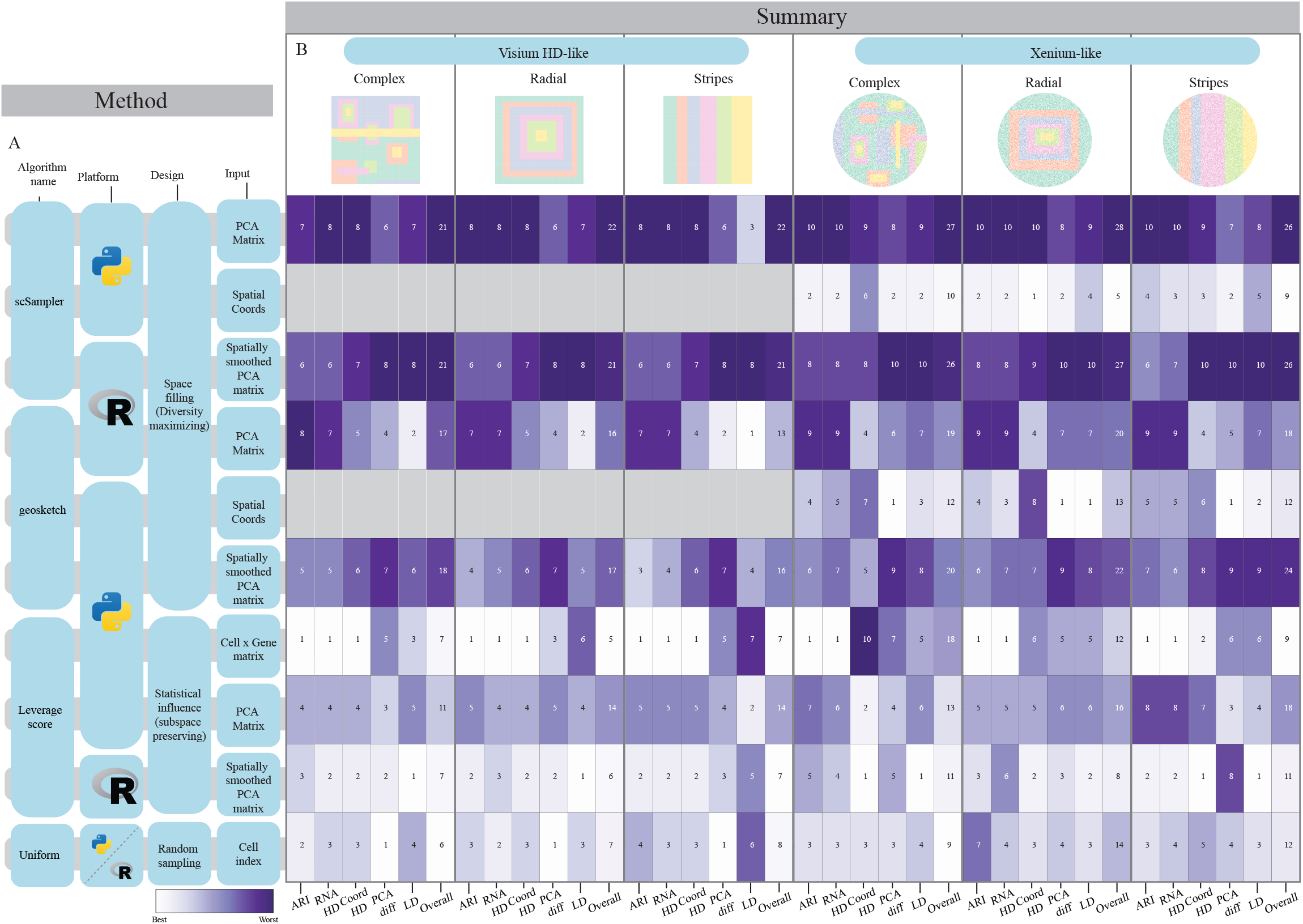
Overall sketching performance for 0.10 sampling fraction across simulated datasets. **A:** Tested sketching algorithm details (platform, sketching approach and input data). **B:** Summary statistics for the six evaluated simulation models. Each column represents a evaluation metric and each row a unique algorithm condition. Numeric values represent the rank order for that metric, color is normalized by column. The final column in each dataset represents the overall spatial score for that dataset, which is the summed ranks of all spatial metrics.

**Figure 4.**
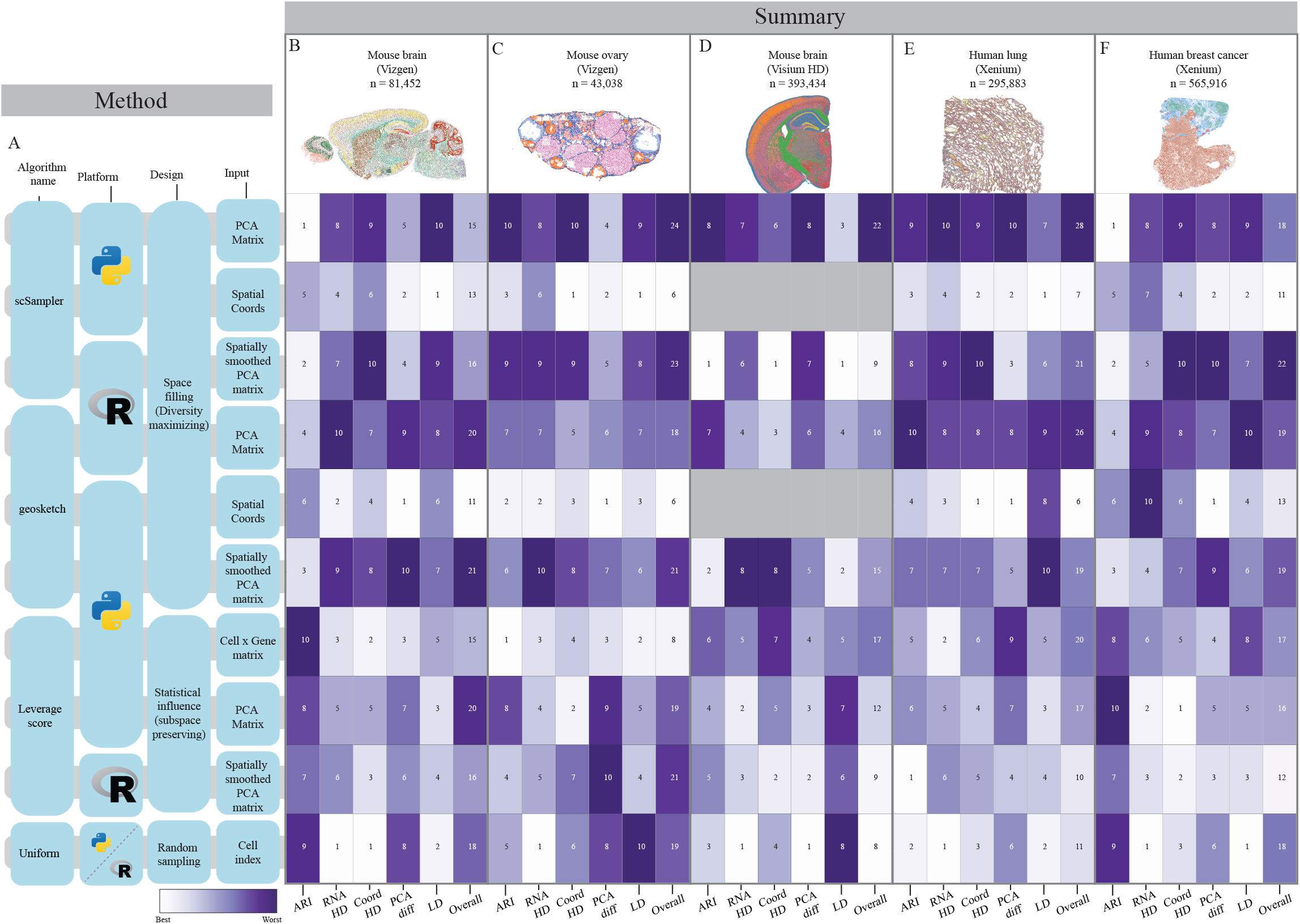
Overall sketching performance for real ST datasets at 0.10 sketching fraction. **A:** Tested sketching algorithm details. **B:** Summary statistics for sagital mouse brain dataset (Vizgen MERFISH). **C:** Summary statistics for mouse ovary dataset. **D:** Summary statistics for coronal mouse brain (10x Visium HD). **E:** Summary statistics for human lung dataset (10x Xenium). **F:** Summary statistics for human breast cancer dataset (10x Xenium). Each column represents a evaluation metric and each row a unique algorithm condition. Numeric values represent the rank order for that metric, color normalized by column. The final column in each dataset represents the overall spatial score for that dataset, which is the summed ranks of all spatial metrics.

### 3.2 Simulated dataset sketching results

This analysis consisted three Visium HD-like and three Xenium-like datasets with different patterns (complex, radial, and striped; see Fig 5A and 5E). For all six datasets, minimization of both transcriptomic Hausdorff distance was similar for all tested algorithms, across sampled fractions, with RNA-based scSampler performing the worst across all datasets, and leverage score performing best (Fig 5B and 5F). For comparison of coordinate-based Hausdorff distances, note that the Visium-like datasets are lacking coordinate-based sketching methods, as they are akin to uniform sampling in this context. For this metric, spatially-smoothed Geosketch and scSampler performed the worst, while uniform and leverage score achieved the smallest Hausdorff distance for the Visium-like dataset, with coordinate-based Geosketch performing best on all the Xenium-like datasets (Fig 5C and 5G). However for both Hausdorff distance comparisons across datasets and between algorithms, all algorithms do a good job minimizing the distance for both coordinates and transcriptomic space, as can been seen with the small variation in values across the sampling fractions. The ARI results show a divergence in algorithm performance across simulated datasets. For the Visium like datasets, there is some variability across sampling fractions, but leverage score and smoothed leverage score, followed by uniform sampling perform best, RNA and smoothed scSampler perform worst (Fig 5D). For the Xenium-like datasets, there is no clear best performing algorithm for the complex and radial datasets, however, scSampler variants still under-perform (Fig 5H). Notably, for both stripe datasets, leverage score and smoothed leverage score significantly outperform all other algorithms (Fig 5D and 5H).

**Figure 5.**
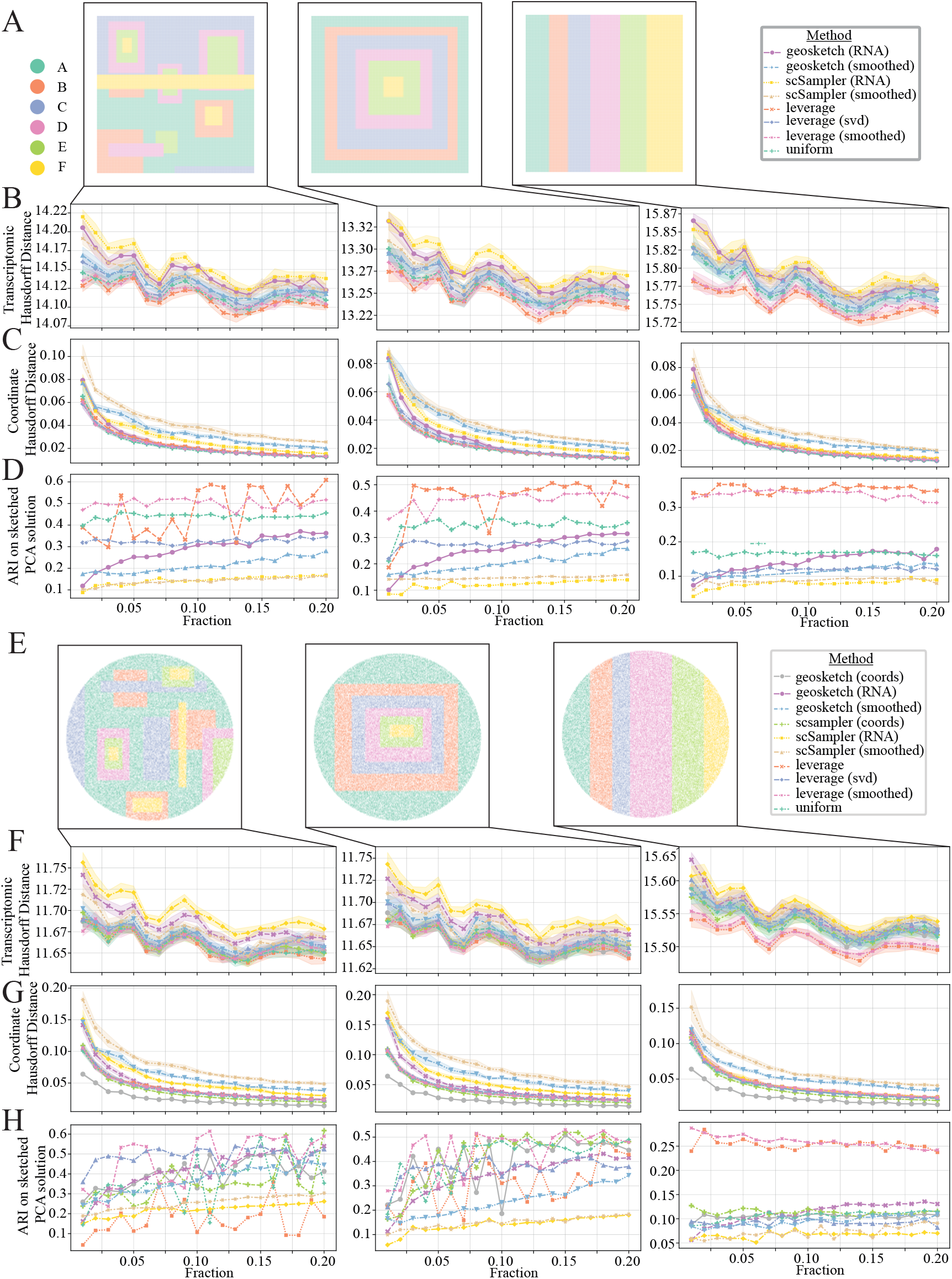
Simulation results for sketching fractions 0.01-0.2. **A:** Ground truth locations plotted spatially for Visium-like complex (left), Visium-like radial (middle) and Visium-like stripes (right) simulated datasets. **B, F:** Transcriptomic Hausdorff distance results for evaluated algorithms for Visium-like datasets (B), and Xenium-like datasets (F). **C, G:** Spatial coordinate Hausdorff distance results for Visium-like datasets (C) and Xenium-like datasets (G). **D, H:** Adjusted Rand Index (ARI) results (sketched clustering results compared to ground truth) for Visium-like datasets (D) and Xenium-like datasets (H).

### 3.3 Human breast cancer

This analysis utilized the 10x Xenium human breast cancer dataset, downloaded through the R package Subcellu-larSpatialData[28]. This resource contains ST data derived from FFPE human breast tissue samples featuring infiltrating ductal carcinoma in situ. Annotations are derived from the H&E stained tissue accompanying the original data (Fig 6A). Transcriptomic sketching results in a higher proportion of cells from the DCIS and normal ducts being sampled (Fig 6B top right), while coordinate-based sketching shows a largely uniform sampling across the tissue, with edge effects (Fig 6B bottom right). Geosketch and scSampler effectively minimize the transcriptomic Hausdorff distance much better than the other methods (Fig 6C top left and Extended data Fig 2). As expected, Coordinate-based sketching effectively minimizes the coordinate Hausdorff distance, with leverage score coming in behind the coordinate-based methods (Fig 6C top right and Extended data Fig 2). Uniform sampling and the coordinate-based Geosketch and scSampler achieved the highest ARI scores for sampling fractions between 0.01 − 0.2, with transcriptomic versions of Geosketch and scSampler under-performing all other methods (Fig 6C middle left). Uniform and coordinate-based sketching methods best preserved local neighborhood structure, with the lowest MSE in location annotation when compared to the full dataset, across all sampled fractions (Fig 6C bottom left and Extended data Fig 2). Areas with the highest MSE for transcriptomic sketching when compared to coordinate-based sketching were localized in areas where transcriptomic sketching tended to over-sample including the DCIS and normal ducts (Fig 6D).

**Figure 6.**
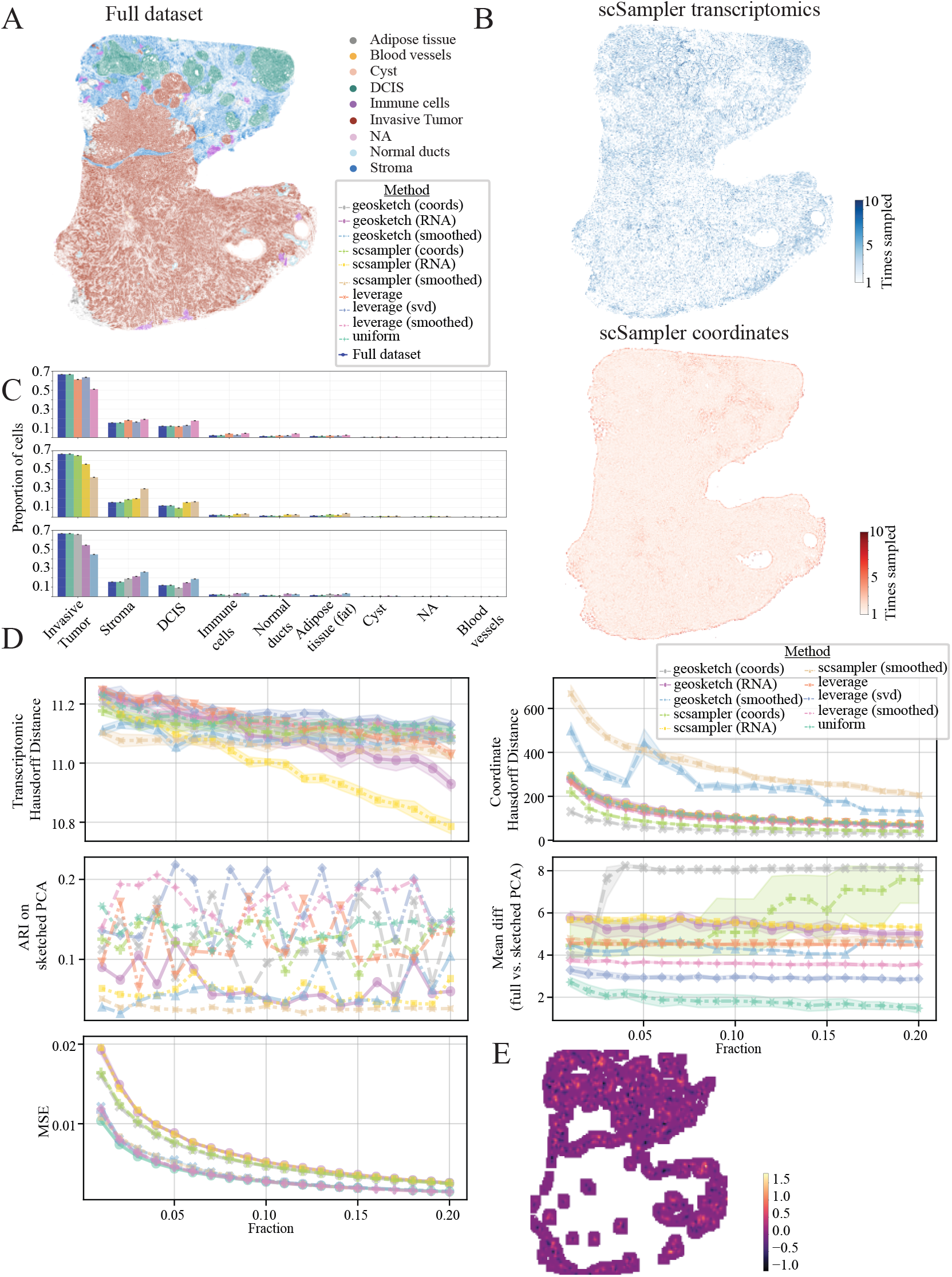
Human breast cancer results (10x Xenium). **A:** Ground truth cell type labels for full dataset **B:** Resulting sketched datasets for transcriptomic (top right), coordinate (bottom right) and composite (bottom left) based scSampler sketching. Color scale indicates how many times a cell was sampled out of ten iterations of the sketching algorithm. **C:** Cell type proportions for full dataset and 0.1 downs-sampled sketches across algorithms evaluated. Rows correspond to Geosketch (top), scSampler (middle) and Leverage score (bottom) algorithms. **D:** Evaluation metrics for sketching algorithms. Transcriptomic Hausdorff distance between sketch and full dataset for sampling fractions from 0.01 − 0.20 (top left), coordinate Hausdorff distance between sketch and full dataset for sampling fractions from 0.01 − 0.20 (top right), Adjusted Rand Index (ARI) for sketched cluster solutions compared to ground truth (middle left), mean difference between the full vs sketched data projected onto the principle component loadings (middle right), mean squared error (MSE) between sketched and full dataset’s local neighborhood composition (bottom left). Colors indicate sketching method. **E:** Difference in local neighborhood composition MSE between transcriptomic vs coordinate-based scSampler sketching at a 10% sampling fraction, visualized on the tissue. Lighter colors indicate higher transcriptomic MSE compared to coordinate based sketching.

### 3.4 Coronal mouse brain results

This analysis was performed on a 10x Visium-HD dataset of a mouse coronal brain section downloaded from the 10x website. Cells were labeled via graph-based clustering as no ground truth annotations were available, and generally correspond to known anatomical structures (Fig 7A). RNA-based sketching significantly over-sampled cells in the hippocampus, pia surface and parts of the thalamus when compared to uniform sampling (Fig 7B). This can also be seen in the quantification of cell type proportions, where we see that both scSampler and Geosketch bias towards label 0 as compared to uniform and leverage scor-based sketching (Fig 7C). All algorithms effectivly minimize the transcriptomic Hausdorff distance (Fig 7D top left), while all variants of leverage score, and uniform sketching best minimize the coordinate Hausdorff distance (Fig 7D top right). Sketched labels compared against graph-based clustering on full dataset (ARI) are highly variable, with no algorithm generating superior performance (Fig 7D middle, left). scSampler, and Geosketch showed similar low MSE in local neighborhood structure preservation compared to the full dataset, while leverage score and uniform sampling had larger values (Fig 7D bottom left).

**Figure 7.**
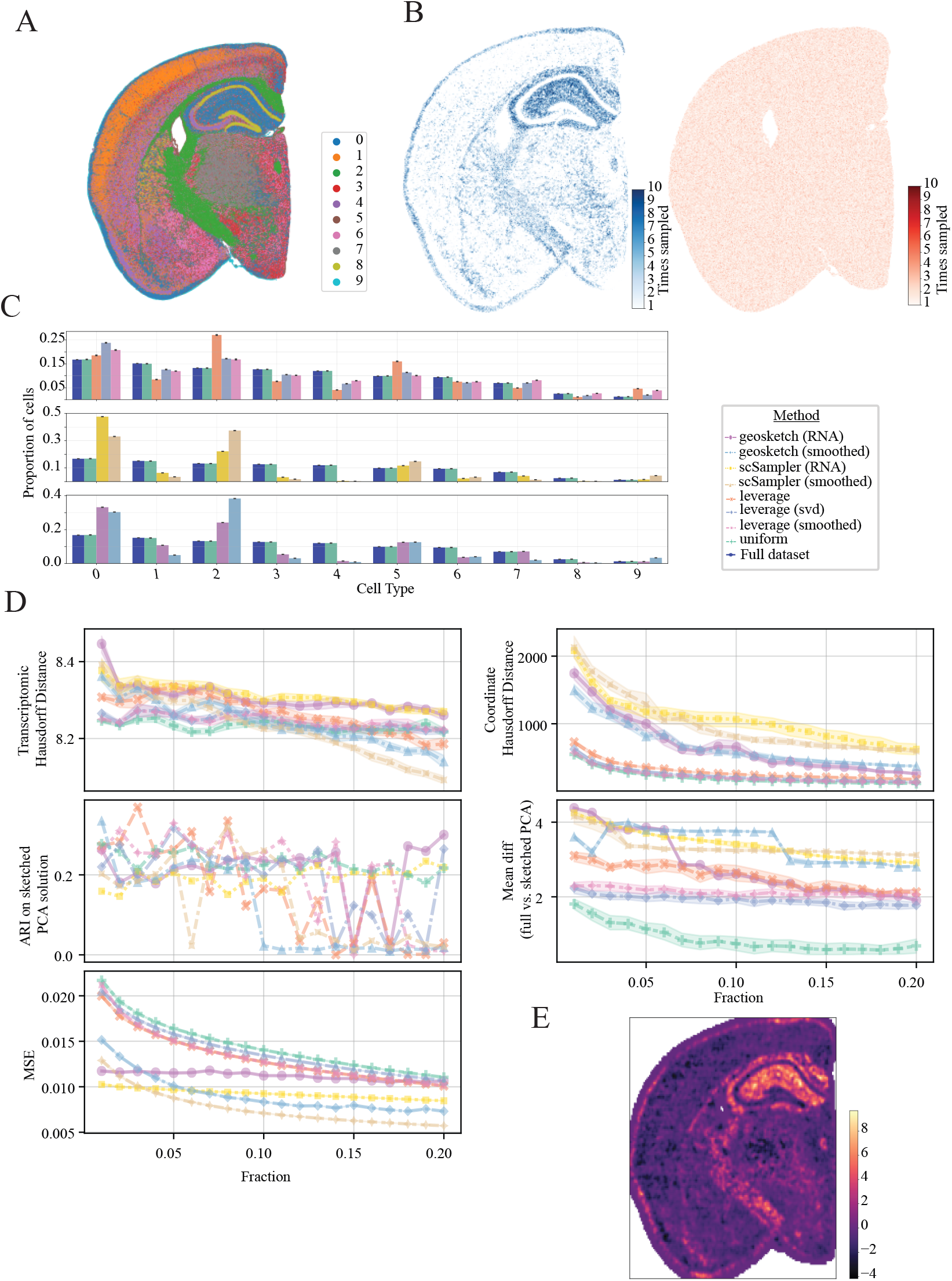
Mouse brain results (10x Visium HD). **A:** Ground truth cell type labels for full dataset. **B:** Sketchedd datasets for scSampler based on transcriptomic data (top right) or tissue coordinates (bottom right). Color scale indicates how many times a cell was sampled out of ten iterations of the sketching algorithm. **C:** Cell type proportions for full dataset and 0.1 down-sampled sketches across algorithms evaluated. Rows correspond to Geosketch (top), scSampler (middle) and Leverage score (bottom) algorithms under different conditions **D:** Evaluation metrics for each sketching algorithm. Transcriptomic Hausdorff distance between sketch and full dataset for sampling fractions from 0.01 − 0.20 (top left), coordinate Hausdorff distance between sketch and full dataset for sampling fractions from 0.01 − 0.20 (top right), ARI for sketched cluster solutions compared to ground truth (middle left), mean difference between the full vs sketched data projected onto the principle component loadings (middle right), MSE between sketched and full dataset’s local neighborhood composition (bottom left). Colors indicate sketching method. **E:** Difference in local neighborhood composition MSE between transcriptomic scSampler vs uniform sketching at a 10% sampling fraction, visualized on the tissue. Lighter colors indicate higher transcriptomic MSE compared to coordinate-based sketching

### 3.5 Human lung results

This analysis utilized the 10x Xenium healthy human lung dataset, downloaded from the 10x website. This resource contains ST data derived from FFPE human lung tissue. Annotations are derived from graph-based Louvain clustering (Extended Data Fig 4A). This fibrotic tissue has a unique heterogeneous structure, making sketching a difficult task. Transcriptomic sketching results in a higher proportion of cells in the bottom left section of the tissue(Extended Data Fig 4B top right), while coordinate-based sketching shows a largely uniform sampling across the tissue, with edge effects (Extended Data Fig 4B bottom right). Interestingly, transcriptomic Geosketch and scSampler have larger transcriptomic Hausdorff distances than the other methods for sampling fractions of 0.01 − 0.2 (Extended Data Fig 4C top left and Extended data Fig 5). Coordinate-based sketching effectively minimizes the coordinate Hausdorff distance, with all other methods comparable (Extended Data Fig 4C top right and Extended data Fig 5). Uniform sampling. the coordinate-based Geosketch and scSampler achieved the highest ARI scores for sampling fractions between 0.01 − 0.2, with transcriptomic versions of Geosketch and scSampler under-performing all other methods (Extended Data Fig 4C middle left). All methods showed similar MSE in local neighborhood structure preservation compared to the full dataset (Extended Data Fig 4C bottom left). This is likely due to the high heterogeneity of the spatial and transcriptomic features of this dataset. Despite this similarity, when comparing between transcriptomic vs coordinate-based scSampler’s MSE it is clear that transcriptomic sketching biases local neighborhoods in areas of higher sampling probability (Extended Data Fig 4D).

### 3.6 Mouse brain results

The mouse brain dataset features detailed cell type classification provided by the Allen Brain Institute [19]. We choose to use a Sagittal section from the atlas as it spans the full anterior-posterior axis and includes both cortical and sub-cortical brain regions, including the cerebellum (Extended Data Fig 6A). Transcriptomic sketching results in a higher proportion of cells in the olfactory bulb and hippocampus being sampled, likely due to a larger amount of transcriptomic variance in this region (Extended Data Fig 6B top right), while coordinate-based sketching shows a largely uniform sampling across the tissue, with pronounced edge effects (Extended Data Fig 6B bottom right). Geosketch and scSampler effectively minimize the transcriptomic Hausdorff distance much better than the other methods (Extended Data Fig 6C top left and Extended data Fig 7). As expected, coordinate-based sketching effectively minimizes the coordinate Hausdorff distance, with leverage score coming in behind the coordinate-based methods, (Extended Data Fig 6C top right and Extended Data Fig 7). Uniform sampling, leverage score, and the coordinate-based Geosketch achieved the highest ARI scores for sampling fractions between 0.01 − 0.2, with the coordinate-based scSampler and transcriptomic versions of Geosketch and scSampler under performing all other methods. This finding highlights the importance of incorporating spatial as well as transcriptomic features into the sketching process (Extended Data Fig 6C middle left). Transcriptomic Geosketch and scSampler under-performed all other methods for local neighborhood structure preservation, with the highest MSE in cell-type proportions when compared to the full dataset, across all sampled fractions (Extended Data Fig 6C bottom left amd Extended Data Fig 7). Areas with the highest MSE for transcriptomic sketching when compared to coordinate-based sketching where localized in areas where transcriptomic sketching tended to over-sample including the olfactory bulb and cerebellum (Extended Data Fig 6D).

### 3.7 Mouse ovary results

This MERFISH dataset features ground-truth cell-level annotations of the mouse ovary and shows clear spatial organization for some cell types (luteal, granulosa) and heterogeneous distribution for others (epithelial, immune, endothelial) (Extended Data Fig 8A)[27]. Transcriptomic sketching results in a higher proportion of cells in the upper left part of the tissue being sampled, likely due to a larger amount of transcriptomic variance in this region (Extended Data Fig 8B top right), while coordinate-based sketching shows a largely uniform sampling across the tissue, with pronounced edge effects**??** (Extended Data Fig 8B bottom right). Transcriptomic Hausdorff distances between the various methods are quite similar, with the leverage score being slightly better across the sampled fractions. Interestingly, coordinate-based sketching achieved a smaller Hausdorff distance compared to the transcriptomic version for both Geosketch and scSampler (Extended Data Fig 8C top left and Extended Data Fig 9). As expected, coordinate-based sketching effectively minimizes the coordinate Hausdorff distance, with leverage score coming in behind the coordinate-based methods (Extended Data Fig 8C top right and Extended Data Fig 9). Coordinate-based sketching and leverage score also achieved the highest ARI scores for sampling fractions between 0.01 − 0.2, with the transcriptomic versions of Geosketch and scSampler under-performing all other methods. This finding highlights the importance of incorporating spatial as well as transcriptomic features into the sketching process (Fig Extended Data Fig 8C middle left). As expected, the coordinate-based sketching best captured the local neighborhood structure, with the lowest MSE in cell-type proportions when compared to the full dataset across all sampled fractions (Extended Data Fig 8C bottom left). Areas with the highest MSE for transcriptomic sketching when compared to coordinate-based sketching were localized in areas where transcriptomic sketching tended to over-sample (Extended Data Fig 8D).

## 4 Discussion

Our benchmarking of sketching algorithms on ST data reveals clear trade-offs between preserving transcriptomic diversity and maintaining tissue coverage. Purely expression-based methods such as Geosketch and scSampler excel at capturing global heterogeneity (i.e., minimizing transcriptional Hausdorff distance and enriching for rare cell states) but do so at the cost of distorting tissue-scale structure. In multiple real tissues (mouse ovary, MERFISH brain, human breast cancer, lung), these methods over-sampled regions of high expression variability (e.g., olfactory bulb, DCIS ducts) and under-sampled more homogeneous areas, leading to biases in downstream neighborhood analyses and local cell-type composition. Conversely, coordinate-only sampling uniformly covers the tissue footprint and achieves minimal coordinate Hausdorff distance, but fails to capture transcriptional extremes and sometimes introduces an edge-effect by over-selecting peripheral cells.

Leverage score methods strike a better balance, particularly when computed on the full cell-by-gene matrix or on a spatially smoothed SVD embedding. Across most datasets and evaluation metrics (i.e., the robust Hausdorff distances, adjusted Rand index of clustering, PCA loading drift, and local cell-type MSE), leverage score-based sketches closely recapitulated both the global transcriptomic landscape and the spatial organization of the tissue. In contrast to pure transcriptomic sketches, spatially-smoothed leverage scores attenuated sampling pattern bias, avoided edge effects, and maintained strong performance on rare cell recovery. Notably, uniform and coordinate sketches performed competitively on datasets with coarsely gridded spots (10x Visium HD and Visium-like simulations), suggesting that when measurement locations are regular and dense, simple schemes suffice.

Our findings underscore the importance of incorporating spatial context into sketch selection for ST studies. While transcriptomic diversity remains critical, especially when rare cell types or subtle expression gradients drive downstream discovery, disregarding spatial structure can lead to misleading inferences about tissue architecture. Spatially-aware sketches, implemented here via a spatial weight matrix and smoothed low-dimensional embeddings, represent a promising compromise: they preserve transcriptomic extremes and spatial uniformity while avoiding unintended edge biases.

Looking forward, these results motivate the development of principled sketching frameworks that explicitly optimize a joint objective over transcriptomic and spatial distances. Possible directions include adaptive weighting of expression versus coordinate-based sampling, multiscale tiling to mitigate grid-edge effects, and integration with biologically informed priors (e.g., known tissue landmarks). Such advances will enable efficient and unbiased down-sampling of ever-larger ST datasets, facilitating rapid exploratory analyses without sacrificing biological fidelity.

## 5 Data Availability

### Mouse ovary

Please contact the authors for access the the Mouse ovary dataset.

### Breast Cancer

We downloaded the Breast cancer 10x Xenium dataset from the **SubcellularSpatialData R** package via **ExperimentHub**, dataset # EH8567, sample ID ’IDC’ (https://www.bioconductor.org/packages/release/data/experiment/html/SubcellularSpatialData.html).

### Human lung

We downloaded the healthy human lung 10x Xenium dataset from publicly available datasets on the 10x website https://www.10xgenomics.com/datasets/xenium-human-lung-preview-data-1-standard.

### Sagittal mouse brain

We downloaded the mouse brain dataset from the Allen Brain Atlas via their **abc atlas access Python** package. Specifically we followed the *MERFISH whole mouse brain STs (Xiaowei Zhuang)* tutorial https://alleninstitute.github.io/abc_atlas_access/notebooks/zhuang_merfish_tutorial.html to access section 3.010 from the ’Zhuang-ABCA-3’ dataset.

### Coronal mouse brain

We downloaded the coronal mouse brain dataset from the publicly available datasets on the 10x website https://www.10xgenomics.com/datasets/visium-hd-cytassist-gene-expression-libraries-of-mouse-brain-he-v4.

### Simulated data

Please contact the authors for access to the simulated datasets.

## 6 Code Availability

See the following Github: https://github.com/gingerii/Benchmarking-sketching-for-spatial-transcriptomics

