## supplemental material for "Benchmarking sketching methods on spatial transcriptomics data"

### Spatial Sketching Supplemental Information

#### List of Algorithms

1 Partial Hausdorff Distance Calculation using Annoy . . . 6
2 Compute PCA projection difference . . . 6
3 local_neighborhood_proportions . . . 7
4 compare_neighborhood_proportions . . . 7

#### List of Tables

1 Summary of simulated datasets and parameters. . . . 3

### 1 Supplemental Methods

#### 1.1 Dataset preprocessing:

##### Breast Cancer (Xenium, 10x):

We downloaded the Breast cancer Xenium dataset from the **SubcellularSpatialData R** package via **ExperimentHub**, dataset # EH8567, sample ID 'IDC' (`https://www.bioconductor.org/packages/release/data/experiment/html/SubcellularSpatialData.html`). Transcripts at each subcellular location were assigned to cells using the `tx2spe` function with `bin = 'cell'`. The resulting `SingleCellExperiment` object was converted to a *Seurat* object, and then converted to an .h5ad file using **SeuratDisk** package's `SaveH5Seurat` and `Convert` functions. Downstream analysis was done in **Python** using the **Scanpy** and **Squidpy** packages. Mitochondrial, ribosomal, and hemoglobin genes identified with the `var_names.str.startswith("MT-"),var_names.str.startswith("RPS","` commands, quality control metrics where calculated with the `calculate_qc_metrics` command, and data were filtered using the `filter_cells` command where `min_genes = 20` and `min_cells = 3`. The filtered data were then log normalized and the top 3000 most highly variable genes where identified using the `highly_variable_genes` function with *favor = 'seurat'*.

##### Coronal mouse brain (Visium HD, 10x)

The Visium HD CytAssist mouse brain (V4) dataset was downloaded from 10x Genomics (`https://www.10xgenomics.com/datasets/visium-hd-cytassist-gene-expression-libraries-of-mouse-brain-he-v4`) and the 8 µm-binned count matrix was loaded in Python via `adata = spatialdata_io.visium_hd(...).data.tables['square_008um'` We annotated mitochondrial genes `adata.var_names.str.startswith("MT-")`, ribosomal genes `adata.var_names.str.s` and hemoglobin genes `adata.var_names.str.match("ĤB[(̂P)]")` and then computed QC metrics with `sc.pp.calculate_` `qc_vars=['mt','rps','rpl','hb'], inplace=True)`. Cells with fewer than 10 detected genes were removed using `adata = filter_cells(adata, min_genes=10)`. Next, counts were normalized to total counts `sc.pp.normalize_tota` log-transformed `sc.pp.log1p(adata)`, and highly variable genes were identified with `sc.pp.highly_variable_genes(adat` (all default parameters).

For dimensionality reduction, we fit a PCA model with 20 components, randomized SVD, and a fixed seed of 2024: `pca_model = PCA(n_components=20, svd_solver='randomized', random_state=2024); pca_data = pca_model.fit_transform(adata.X.toarray()); adata.obsm['X_pca'] = pca_data`. Finally, we constructed a neighborhood graph (`sc.pp.neighbors(adata)`), computed a UMAP embedding (`sc.tl.umap(adata)`), and performed community detection via Leiden (`sc.tl.leiden(adata)`) and Louvain (`sc.tl.louvain(adata)`). All functions were executed with default settings unless otherwise specified.

##### Healthy lung (Xenium, 10x):

The Xenium human lung preview dataset was obtained from 10x Genomics (`https://www.10xgenomics.com/datasets/xenium-human-lung-preview-data-1-standard` ) and loaded into an AnnData object. Quality control metrics were computed with `sc.pp.calculate_qc_metrics(adata, percent_top=(10, 20, 50, 150), inplace=Tr` after which cells with fewer than 10 total counts were removed via `sc.pp.filter_cells(adata, min_counts=10)` and genes detected in fewer than 5 cells were discarded using `sc.pp.filter_genes(adata, min_cells=5)`.

The filtered counts were normalized to total counts per cell (`sc.pp.normalize_total(adata, inplace=True)`), log-transformed (`sc.pp.log1p(adata)`), and reduced to 20 principal components (`sc.pp.pca(adata)`). A neighborhood graph was then constructed (`sc.pp.neighbors(adata)`), followed by UMAP embedding (`sc.tl.umap(adata)`), Leiden clustering (`sc.tl.leiden(adata)`), and Louvain clustering (`sc.tl.louvain(adata)`). All functions were run with default parameters unless otherwise noted.

##### Sagittal mouse brain (MERFISH, Vizgen):

The MERFISH whole mouse brain spatial transcriptomics dataset (Zhuang-ABCA-3, section 3.010) was accessed via the `abc_atlas_access` Python package following the "MERFISH whole mouse brain spatial transcriptomics (Xiaowei Zhuang)" tutorial (`https://alleninstitute.github.io/abc_atlas_access/notebooks/`

zhuang_merfish_tutorial.html ). The selected slice was loaded into an AnnData object (adata). Quality control metrics were calculated with sc.pp.calculate_qc_metrics(adata, percent_top=(10,20,50,150), inplace=True), after which cells with fewer than 10 total counts were removed via sc.pp.filter_cells(adata, min_counts=10) and genes detected in fewer than 5 cells were discarded using sc.pp.filter_genes(adata, min_cells=5). The filtered count matrix was then normalized to total counts per cell (sc.pp.normalize_total(adata, inplace=True)), log-transformed (sc.pp.log1p(adata)), and reduced by PCA (sc.pp.pca(adata)). Finally, we constructed a neighborhood graph (sc.pp.neighbors(adata)), computed a UMAP embedding (sc.tl.umap(adata)), and performed Leiden (sc.tl.leiden(adata)) and Louvain (sc.tl.louvain(adata)) clustering. All functions were run with default parameters unless otherwise specified.

**Mouse Ovary:**

The dataset was provided directly by the authors [1] and was already preprocessed. RASP was applied directly to the AnnData object provided. Briefly, processing included cell segmentation (CellPose and MERlin)[2, 3] for the acquisition of cell and transcript data. Cells with fewer than 10 transcript counts were excluded from further analysis to ensure data quality. Data was log-normalized, scaled, PCA was performed, neighborhood identification, and cell clusterings calculated. To determine the cell type for each cluster, transcript counts were compiled for each gene across clusters, focusing on known markers within the top 10 identified transcripts. Differential expression analysis was conducted using **Seurat** v3 in **R**, allowing the identification of specific markers associated with each Leiden cluster. Additionally, visualization of spatial regions was achieved using **Squidpy** [4] in conjunction with the **AnnData** [5] and **Scanpy** [6] libraries.

**Simulated data:**

Raw simulated count data were processed in **Python** using the **Scanpy** and **Squidpy** packages. Count matrices were normalized to medial total counts using the pp.normalize_total command, logarithmized with the pp.log1p command and variable genes identified using the pp.highly_variable_genes function, all with default parameters. See Table 1 for simulation parameters.

Table 1: Summary of simulated datasets and parameters.

| Item | Description |
|---|---|
| Platforms | Visium-like (grid), Xenium-like (random in rectangle) |
| Spatial layouts | Complex (2), Radial (2), Striped (2) |
| Datasets | 6 total (3 Visium-like, 3 Xenium-like) |
| Size per dataset | 100,000 locations, 500 genes |
| Labels | 6 classes (A–F) |
| Simulation tool | SRTsim (R), seed = 1 |
| Gene composition | 300 high-signal, 150 low-signal, 50 noise |
| Dispersion | 0.5 |
| Baseline mean | 1 |
| Zero-inflation | 0.8 |
| Class A LFC | 1 |
| Class B LFC | 2 |
| Class C LFC | 4 |
| Class D LFC | 6 |
| Class E LFC | 3 |
| Class F LFC | 12 |

#### 1.2 Evaluated sketching algorithms

- **Uniform sampling:** Sketched cells where randomly selected from the *adata* object's indices without replacement using uniform probabilities of $1/n$ . Indices were used to subset the full dataset to create the sketch. Each iteration was performed using a different seed to generate distinct sketches for each experiment.

- **Leverage score:**
  - **Cell-by-gene matrix version:** The full dataset leverage scores were calculated using **Seurat's** *LeverageScore* function in R with default parameters. Seurat requires the *FindVariableFeatures* function to be executed prior to running the leverage score calculation. For probe-based ST datasets (cancer, lung, sagittal brain, ovary, simulated datasets) we set the *nfeatures* argument to the total number of genes measured in the dataset so as to not lose any information. For the coronal brain dataset, *nfeatures* was set to 2000.
  - **SVD version:** To quantify the influence (leverage) of each sample (cell) on the low-rank approximation of our data, we proceeded as follows. Let $X \in R^{n \times p}$ be the full data matrix of $n$ samples by $p$ features. We used the randomized truncated singular value decomposition (SVD) implementation from scikit-learn to obtain the $k$ leading singular vectors:

    $$X \approx U_k \, S_k \, V_k^T, \quad U_k \in R^{n \times k}, \; S_k \in R^{k \times k}, \; V_k \in R^{p \times k}.$$

    We set $k = 20$ and fixed the random seed to 42 for reproducibility. Once $U_k$ is obtained, the leverage score of sample $i$ is defined by

    $$\ell_i \;=\; \sum_{j=1}^{k} \bigl(U_k\bigr)_{ij}^2 \, .$$
  - **smoothed SVD version:**

  Scores were saved as .csv files and added to the *adata.obs* slot in Python. The scores where then normalized to the total sum to create sampling probabilities, and sketches where selected without replacement using the probabilities derived from the leverage scores.

- **Geometric sketching:**
  - **Transcriptomic version:** Geometric sketching approximately minimizes the Hausdorff distance between the sub-sampled and full datasets utilizing a minimax distance design [7] with the goal of evenly sampling across the transcriptomic space. The transcriptomic data was first reduced to 20 PCs. The resulting matrix was then passed to **Geosketch's** *gs* function which returns indices of the sketch.
  - **Coordinate version:** For coordinate-based sketching, the coordinate matrix was passed directly to the *gs* function.

- **scSampler sketching:**
  - **Transcriptomic version:** scSampler also seeks to minimize the Hausdorff distance between the sub-sampled and full datasets, but, unlike geometric sketching, employs the maximin distance design to make cells in the subsample as separable as possible. scSampler has been shown to outperform other methods in terms of minimizing the Hausdorff distance [8]. For transcriptomic-based sketching, the ST data was first reduced to 20 PCs, and the resulting matrix was passed to the *scsampler* function with *random_split* = 16 to speed up the computation.
  - **Coordinate version:** For coordinate-based sketching, the coordinate matrix was passed directly to the *scsampler* function.

- **Spatially smoothed versions of scSampler, Geosketch, Leverage SVD:**
  - We first constructed a spatial weights matrix $W \in R^{n \times n}$ from the spatial coordinates stored in `adata` using the RASP Python package:

    ```
    weights = RASP.build_weights_matrix(
        adata,
        n_neighbors = kNN_threshold,
        beta        = beta,
        platform    = platform
    )
    ```

- The un-smoothed low-dimensional embedding $U$ (for Leverage SVD: the truncated left singular vectors; for Geosketch and scSampler: the PCA score matrix) was then spatially smoothed by left-multiplication with $W$:

$$\widetilde{U} = W U,$$

  implemented in Python as:

```
smoothed_U = weights @ csr_matrix(U)
```

- The smoothed embedding $\widetilde{U}$ was used in place of $U$ to:
  * re-compute leverage scores (Leverage SVD),
  * pass into `Geosketch::gs` (Geosketch),
  * or pass into `scsampler` (scSampler).
- The RASP parameters `kNN_threshold` and $\beta$ were set, per dataset, to the recommended values:
  * Mouse ovary (MERFISH): $kNN = 5,\ \beta = 2$
  * Allen sagittal mouse brain (MERFISH): $kNN = 20,\ \beta = 2$
  * Simulation datasets: $kNN = 30,\ \beta = 0$
  * 10x Visium HD brain: $kNN = 30,\ \beta = 2$
  * 10x Xenium lung: $kNN = 10,\ \beta = 2$
  * 10x Xenium cancer: $kNN = 60,\ \beta = 0$

#### 1.3 Evaluation metrics

##### 1.3.1 Hausdorff Distance calculation

We calculated the Hausdorff distance between the cell-by-gene matrix as well as the spatial coordinates for the sub-sampled dataset and the original dataset. This metric measures the maximum distance between two sets of points, serving as an indicator of the differences in the expression profiles and spatial distribution between the subsample and original. The exact Hausdorff distance is sensitive to outliers, so for evaluation we computed the partial or robust Hausdorff distance. Also note that we used Annoy distance estimation to speed up distance calculations (see SI X.X). The robust Hausdorff distance is calculated as follows:

##### 1.3.2 Adjusted Rand Index (ARI) calculation

We compared the sketched clustering results against the ground truth labels using the ARI[9] as implemented by the `adjusted_rand_score` function from the **sklearn** package (version 1.5.2). Mathematically, the ARI is defined as:

$$\text{ARI} = \frac{\sum_{ij}\binom{n_{ij}}{2} - \left[\sum_i \binom{a_i}{2}\sum_j \binom{b_j}{2}\right]/\binom{n}{2}}{\frac{1}{2}\left[\sum_i \binom{a_i}{2} + \sum_j \binom{b_j}{2}\right] - \left[\sum_i \binom{a_i}{2}\sum_j \binom{b_j}{2}\right]/\binom{n}{2}}$$

Where:

- $n$ is the total number of elements (e.g., cells in spatial domains or annotations).
- $n_{ij}$ is the number of elements that are in cluster $i$ in the first partition and in cluster $j$ in the second partition.
- $a_i$ is the number of elements in cluster $i$ of the first partition.
- $b_j$ is the number of elements in cluster $j$ of the second partition.
- $\binom{n}{2}$ is the binomial coefficient, representing the number of ways to choose 2 elements from $n$, calculated as:

$$\binom{n}{2} = \frac{n(n-1)}{2}$$

The ARI formula adjusts for the chance similarity between clusters by considering both pairwise agreements and disagreements. The numerator counts the agreements, and the denominator normalizes the score, yielding an index that ranges between -1 (no agreement) and 1 (perfect agreement), with 0 indicating random labeling.

**Algorithm 1** Partial Hausdorff Distance Calculation using Annoy

Inputs:

- $array1$: Array of points $(n, d)$
- $array2$: Array of points $(m, d)$
- $q$: float, optional, between 0 and 1
- $metric$: string, optional, distance metric ('euclidean' or 'angular')
- $n\_trees$: int, optional, number of trees for Annoy distance estimation. See SI X.X for details on Annoy estimation.

Outputs:

- Estimated partial Hausdorff distance

Notation:

- $w$: Current weights (parameters)
- $distances$: List of distances computed

1: $distances \leftarrow$ Compute_Distances_Annoy$(array1, array2, n_trees)$ ▷ Compute distances from array1 to array2 using Annoy estimation
2: Sort $distances$ ▷ Sort the computed distances to prepare for finding the Kth largest value
3: $K \leftarrow$ Floor$((1 - q) \times$ Length$(distances))$ ▷ Determine the index of the Kth largest value based on the quantile parameter q
4: if $K > 0$ then return $distances[K - 1]$ ▷ Return the Kth largest distance if K is greater than 0
5: elsereturn $distances[0]$ ▷ If K is 0 or less, return the smallest distance

##### 1.3.3 PCA distance calculation

To assess whether the sketched datasets capture the structure and variability of the full dataset, we computed the difference between the projection of the sketched data onto the full dataset PCs and sketched dataset PCs. The calculation was done as follows:

**Algorithm 2** Compute PCA projection difference

Inputs:

- $\mathbf{F}$: Full data matrix of shape $(m \times d)$
- $\mathbf{S}$: Sketched data matrix of shape $(n \times d)$
- $k$: Number of principal components to compute

Outputs:

- Mean distance between PC projections

1: $\mathbf{W}_{full} \leftarrow$ PCA$(\mathbf{F}, n_{\texttt{components}})$ ▷ Compute the top $k$ PC loading vectors for the full data
2: $\mathbf{W}_{sketch} \leftarrow$ PCA$(\mathbf{S}, n_{\texttt{components}})$ ▷ Compute the top $k$ PC loading vectors for the sampled data
3: $\mathbf{P}_f \leftarrow \mathbf{S}\mathbf{W}_{full}$ ▷ Project sketched data onto full data PCs
4: $\mathbf{P}_s \leftarrow \mathbf{S}\mathbf{W}_{sketch}$ ▷ Project sketched data onto the sketched data PCs
5: $distances \leftarrow$ euclidean_distance$(P_f, P_s)$ ▷ Compute distances
6: $mean\_distance \leftarrow$ mean$(distances)$
return $mean\_distance$

##### 1.3.4 Local neighborhood difference analysis

We conducted local spatial neighborhood analysis on the full and sketched datasets to investigate which method best preserves local cell type proportions. To do this we calculated the proportion of cell types for kNN away from each cell in the sketched and full datasets, and computed the mean squared error (MSE) between the two. See Algorithm 3 and 4 for details.

**Algorithm 3** local_neighborhood_proportions

Inputs:

- $\mathbf{L_{possible}}$: Array of possible labels from the full dataset
- $\mathbf{L}$: Array of cell type or domain labels from the dataset
- $\mathbf{C}$: Array of XY coordinates from dataset
- $k$: Number of nearest neighbors to consider
- *Annoy_index*: Index of approximate nearest neighbor distances and index pointers. See SI X.X for details on construction.

Outputs:

- Cell type proportions matrix

```
type_index_dict ← {}                      ▷ Initialize empty data structure for keeping track of cell types
for i, t ∈ enumerate(L_possible) do                                   ▷ iterate over enumerated list
    type_index_dict[t] ← i                     ▷ Assign the cell type 't' to its corresponding index 'i'
end
n_cells, n_dims ← shape(C)                                      ▷ Get number of cells and dimensions
proportions ← []                                               ▷ Initialize empty array of proportions
for i ∈ {n_cells} do
    neighbors ← Annoy_index[i, k][1 :]  ▷ Use the annoy index to get extract the 'k' neighbor indices for a
given cell, minus itself
    neighbor_type ← L[neighbors]            ▷ Get the cell type labels of the neighbors for given cell
    for t ∈ {neighbor_type} do
        proportions[i, type_index_dict[t]+ = 1      ▷ Count occurrences of cell types 't' around given cell 'i'
    end
    proportions[i] = /k                                                      ▷ Normalize to proportions
end
    return proportions
```

**Algorithm 4** compare_neighborhood_proportions

Inputs:

- $\mathbf{M_1}$: Cell type proportion matrix from full dataset
- $\mathbf{M_2}$: Second cell type proportion matrix from sketched dataset

Outputs:

- $MSE$: Mean squared error between the two matrices (for cells contained in sketched matrix)

```
sketch_index ← M2.index                  ▷ Get indices for the sketched data cell type proportion matrix
m1 ← M1[sketch_index]                                       ▷ Subset the M2 matrix by sketched indices
MSE ← MSE(M2, m1)                                  ▷ Compute mean squared error between the two matrices
    return MSE
```

##### 1.3.5 Overall spatial score and comparison

To summarize all algorithm evaluations across tested datasets, we calculated individual metric ranks for ARI, transcriptomic and coordinate Hausdorff distance, difference in PCA matrices, and MSE of the local neighborhood distortion for a single sampling fraction of 0.1, as visualized in Figs **??**, **??**. For the ARI metric, rank was ordered from highest to lowest value, for the other metrics, rank was ordered from lowest to highest value. The "Overall" rank is simply the aggregated rank sum for the spatially relevant metrics (ARI, coordinate Hausdorff distance and local distortion).

##### 1.3.6 A note on k-Nearest Neighbor (kNN) lookup

Our analysis required computing kNN distances frequently, for both the local neighborhood analysis and calculating the Hausdorff distance. To improve computational speed across our experiments we utilized the **Annoy** (version 1.17.2) C++ package with Python bindings. We calculated approximate Euclidean distances between the coordinates of our ST datasets using the *AnnoyIndex*, *add_items*, and *build* functions, with $n\_trees = 20$.

#### 1.4 A note on edge effects

Coordinate-based sketching using Geosketch and scSampler over-samples cells on the edge of the tissue. Both algorithms tile or overlay a grid on the tissue. Interior grid-cells are packed full of points, so "one point per cell" over-samples high-density regions unless you down-weight them. On the very edge of the tissue, the grid-cells lie only partially over data, and contain fewer points. The net result is a larger fraction of picks coming from those partial/low-density edge cells. Cells on the edge have larger empty neighborhoods and so are deemed more "informative" simply because no other points sit beyond them in that direction.

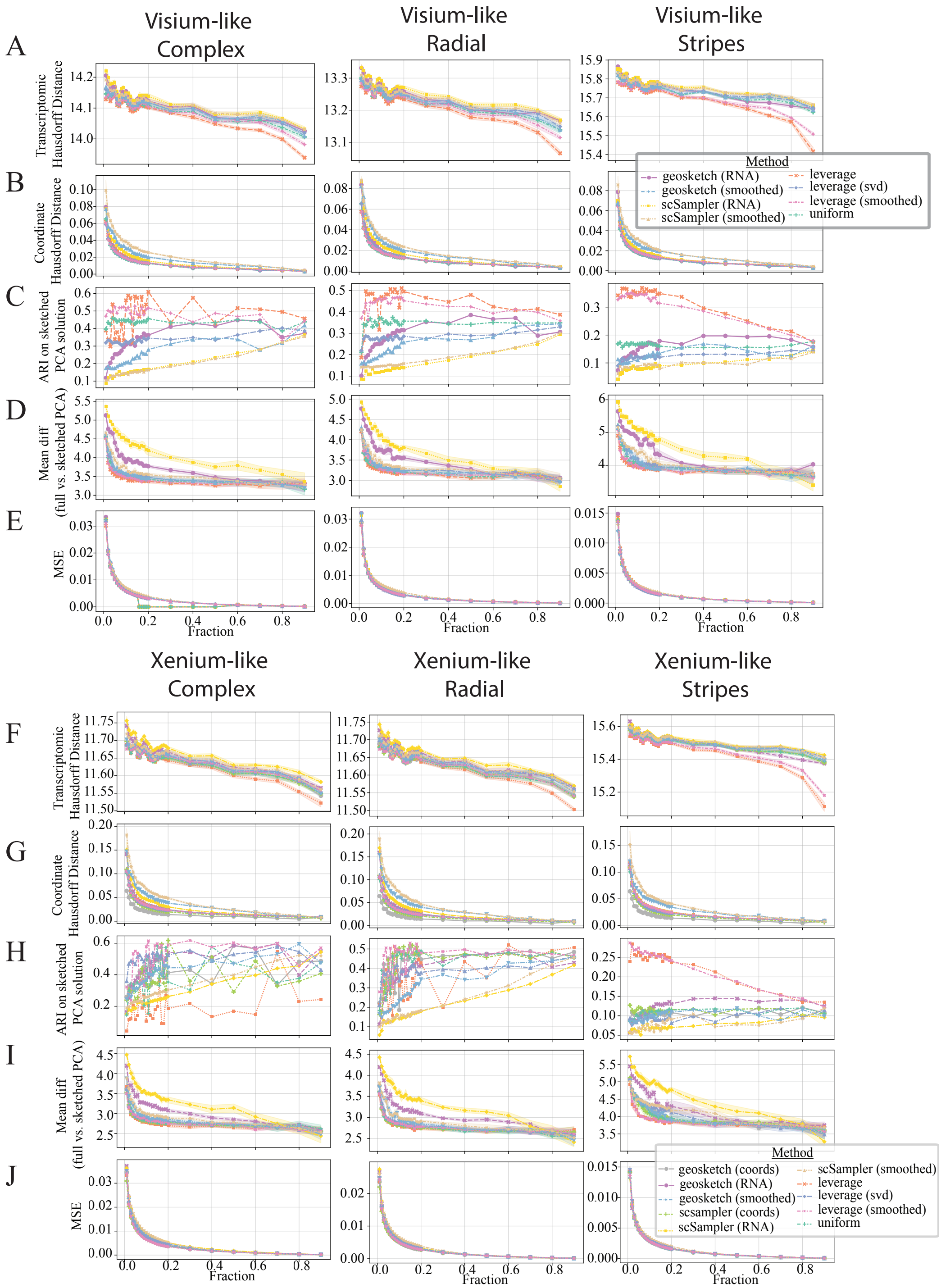

Extended Data Fig. 1: **Sketching evaluation for sketching fractions 0.01-0.9.** **A, F:** Transcriptomic Hausdorff distance for visium-like (A) and xenium-like datasets (F). **B, G:** Coordinate Hausdorff distance for visium-like (B) and xenium-like datasets (G). **C, H:** ARI for visium-like (C) and xenium-like (H) datasets. **D, I:** Mean difference between full and sketched PCA matrices for visium-like (D) and xenium-like (I) datasets. **E, J:** Mean squared error in local neighborhood composition (kNN = 10) for visium-like (E) and xenium-like (J) datasets.

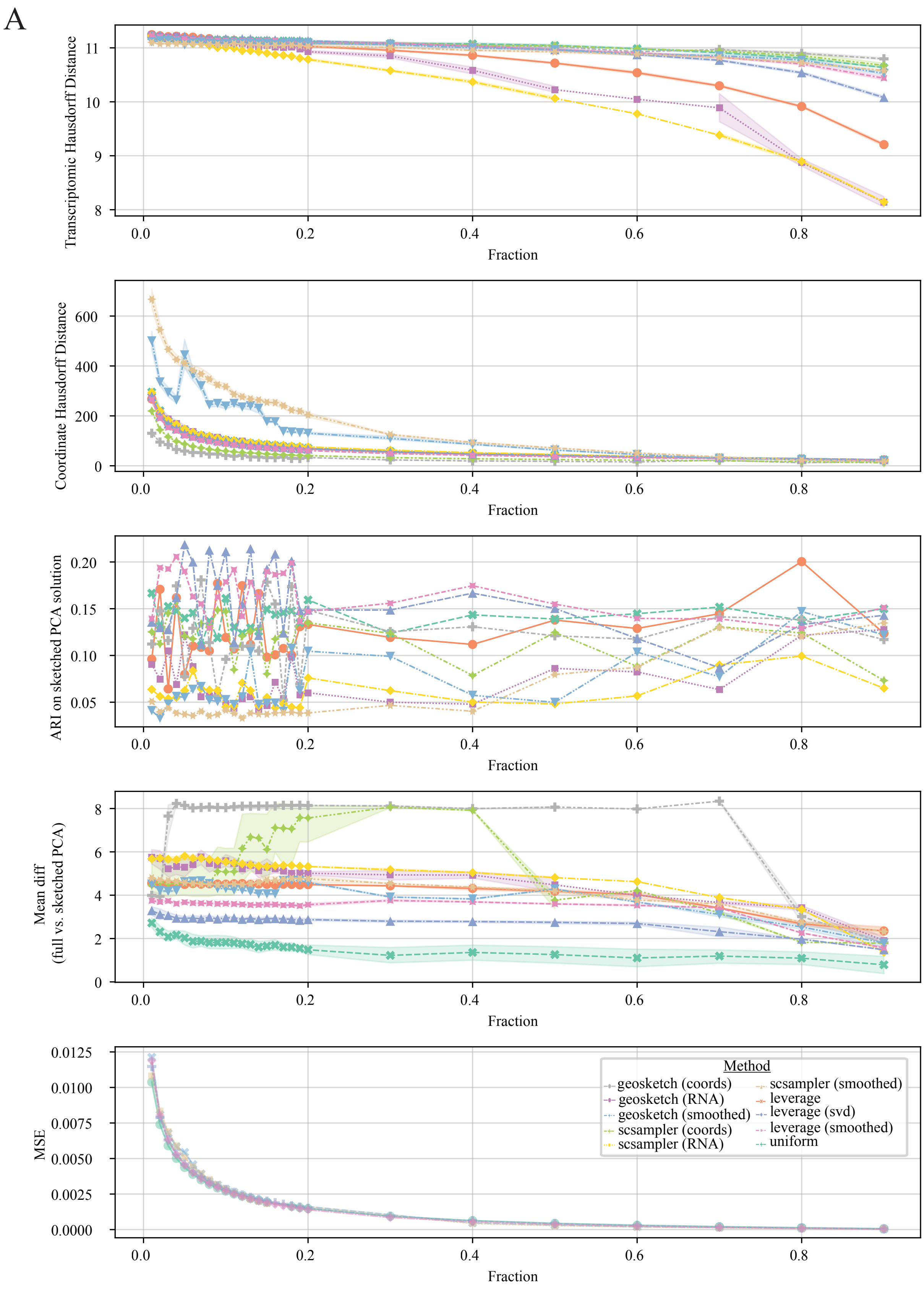

Extended Data Fig. 2: **Sketching evaluation across sample fractions 0.01-0.9 for the human breast cancer dataset.** **A:** From top to bottom, metrics shown are Transcriptomic Hausdorff distance, coordinate Hausdorff distance, ARI, Mean difference between full and sketched PCA matrices, and mean squared error in local neighborhood composition (kNN = 10).

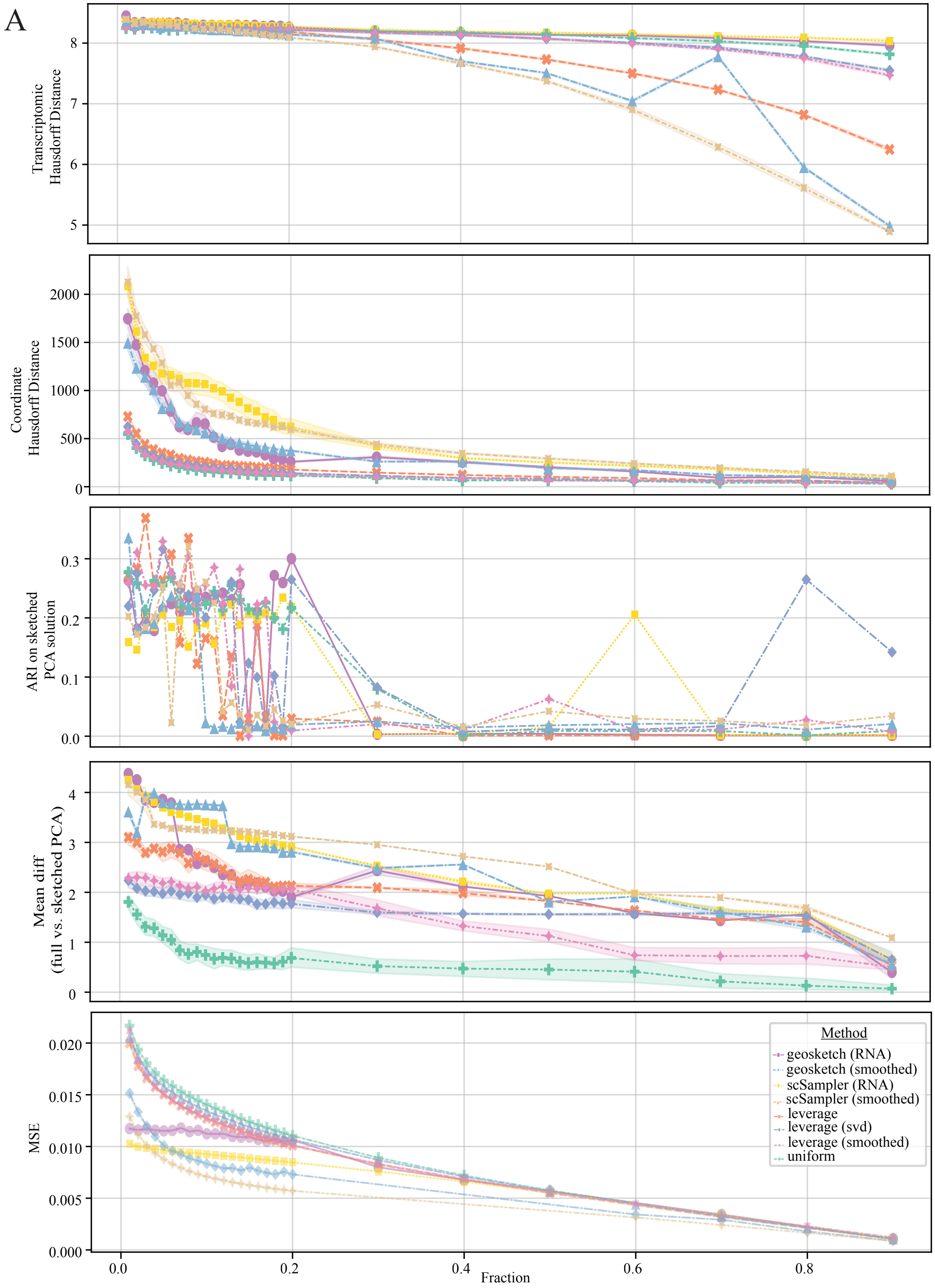

Extended Data Fig. 3: **Sketching evaluation across sample fractions 0.01-0.9 for the coronal mouse brain (Visium HD) dataset. A:** From top to bottom, metrics shown are Transcriptomic Hausdorff distance, coordinate Hausdorff distance, ARI, Mean difference between full and sketched PCA matrices, and mean squared error in local neighborhood composition (kNN = 10).

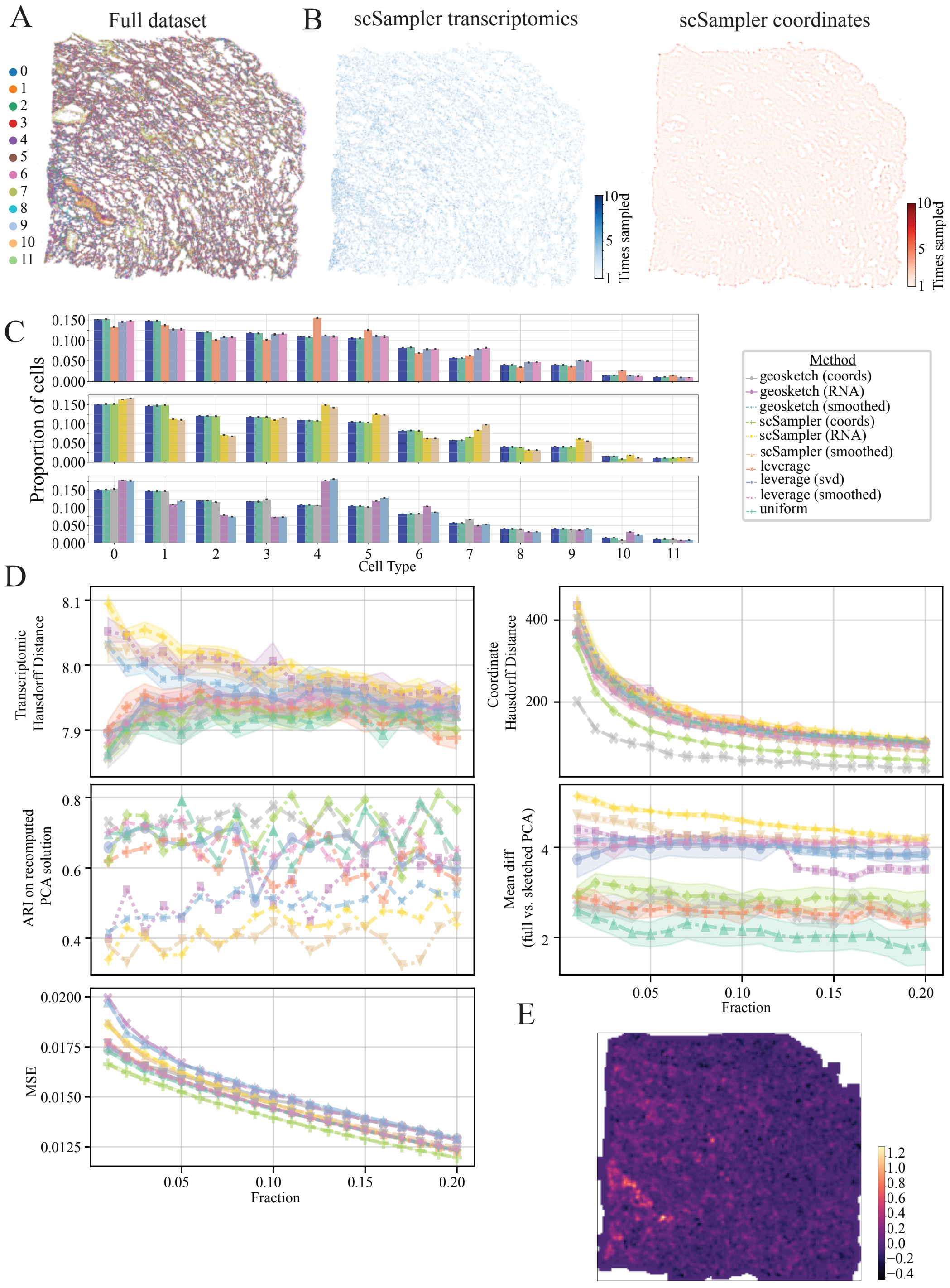

Extended Data Fig. 4: **Human lung (10x Xenium). A:** Ground truth cell type labels for full dataset **B:** Resulting sketched datasets for transcriptomic (top right), coordinate (bottom right) and composite (bottom left) based scSampler sketching. Color scale indicates how many times a cell was sampled out of ten iterations of the sketching algorithm **C:** Cell type proportions for full dataset and 0.1 down-sampled sketches across algorithms evaluated. **D:** Evaluation metrics for sketching algorithms. Transcriptomic Hausdorff distance between sketch and full dataset for sampling fractions from $0.01-0.20$ (top left), coordinate Hausdorff distance between sketch and full dataset for sampling fractions from $0.01-0.20$ (top right), Adjusted rand index score for sketched cluster solutions compared to ground truth (middle left), mean difference between the full vs sketched data projected onto the principle component loadings (middle right), mean squared error (MSE) between sketched and full dataset's local neighborhood composition. Colors indicate sketching method. **E:** Difference in local neighborhood composition MSE between transcriptomic vs coordinate based scSampler sketching at a 10% sampling fraction, visualized on the tissue. Lighter colors indicate higher transcriptomic MSE compared to coordinate based sketching

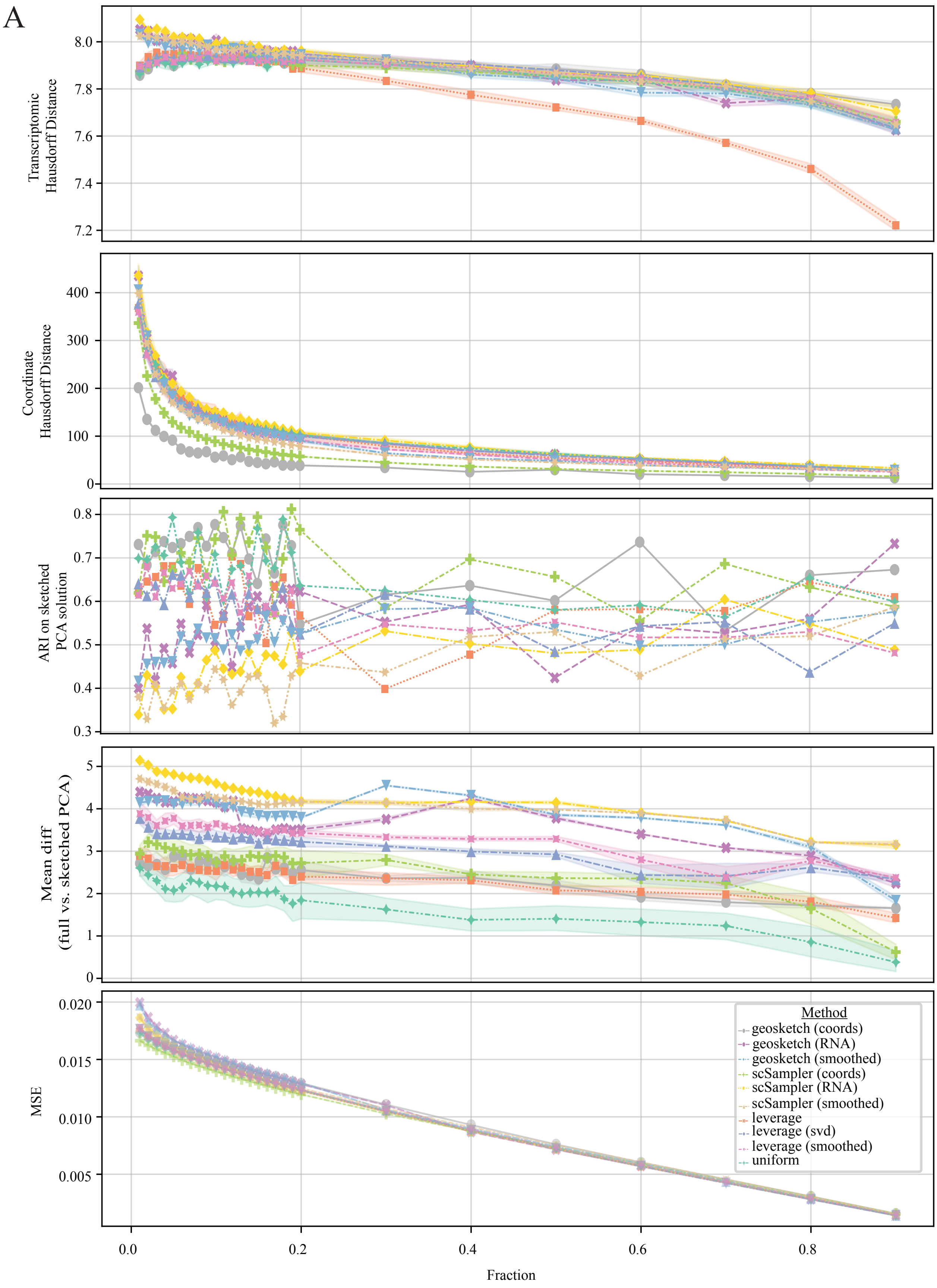

Extended Data Fig. 5: **Sketching evaluation across sample fractions 0.01-0.9 for the human lung dataset. A:** From top to bottom, metrics shown are Transcriptomic Hausdorff distance, coordinate Hausdorff distance, ARI, Mean difference between full and sketched PCA matrices, and mean squared error in local neighborhood composition (kNN = 10).

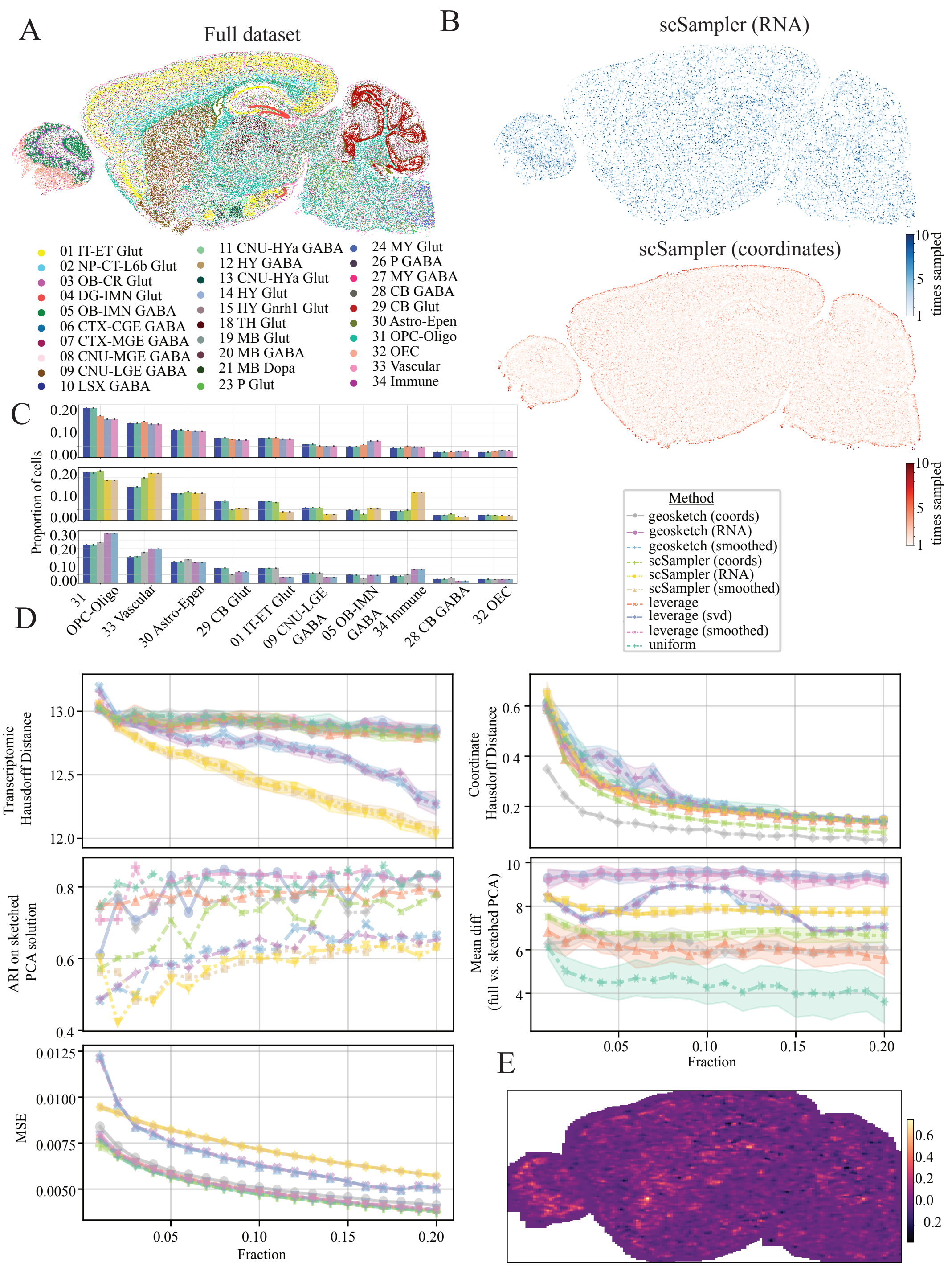

Extended Data Fig. 6: **Mouse brain results (Vizgen MERFISH). A:** Ground truth cell type labels for full dataset **B:** Resulting sketched datasets for transcriptomic (top right), coordinate (bottom right) and composite (bottom left) based scSampler sketching. Color scale indicates how many times a cell was sampled out of ten iterations of the sketching algorithm **C:** Cell type proportions for full dataset and 0.1 down-sampled sketches across algorithms evaluated. **D:** Evaluation metrics for sketching algorithms. Transcriptomic Hausdorff distance between sketch and full dataset for sampling fractions from $0.01 - 0.20$ (top left), coordinate Hausdorff distance between sketch and full dataset for sampling fractions from $0.01 - 0.20$ (top right), Adjusted rand index score for sketched cluster solutions compared to ground truth (middle left), mean difference between the full vs sketched data projected onto the principle component loadings (middle right), mean squared error (MSE) between sketched and full dataset's local neighborhood composition. Colors indicate sketching method. **E:** Difference in local neighborhood composition MSE between transcriptomic vs coordinate based scSampler sketching at a 10% sampling fraction, visualized on the tissue. Lighter colors indicate higher transcriptomic MSE compared to coordinate based sketching

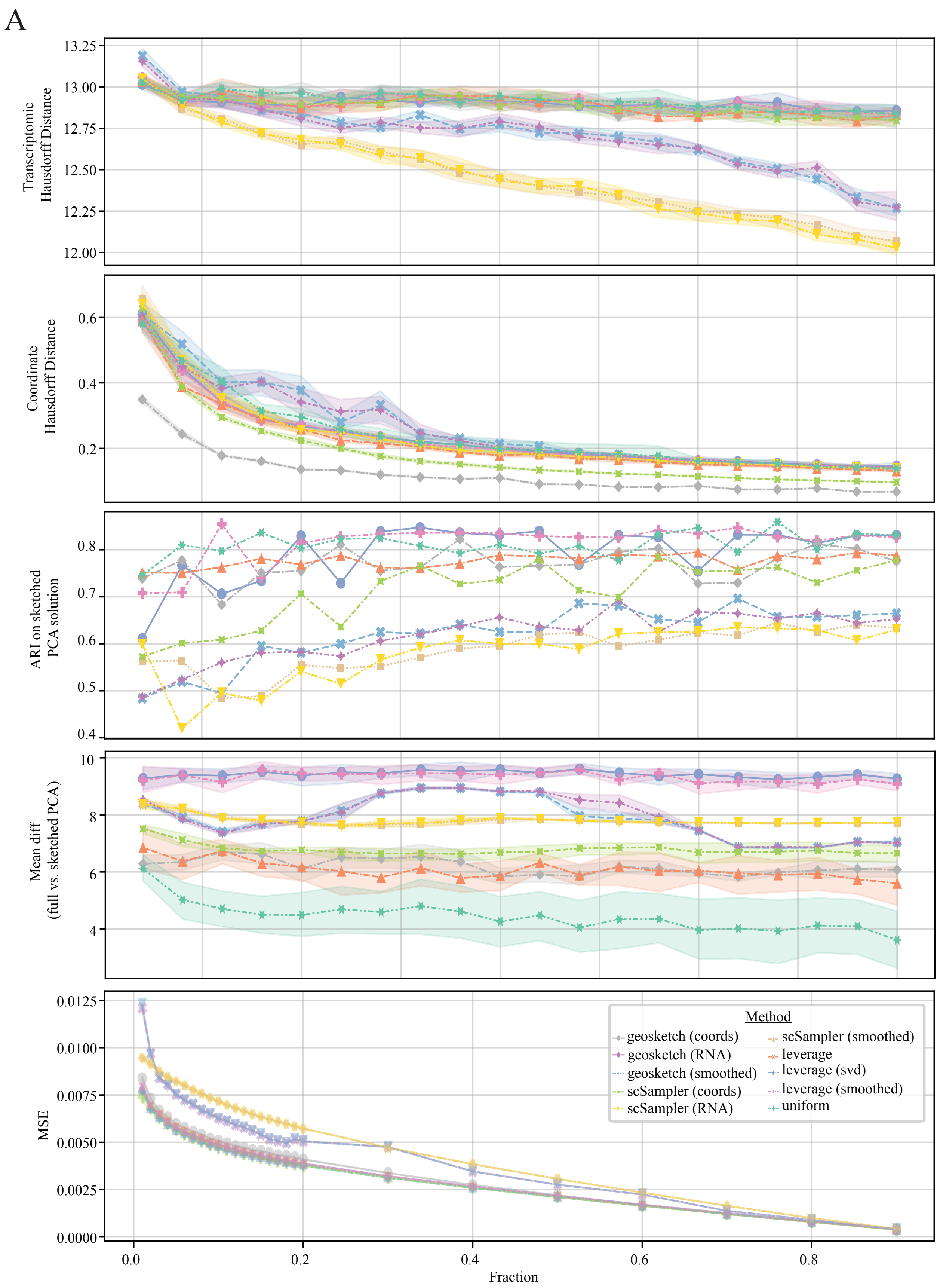

Extended Data Fig. 7: **Sketching evaluation across sample fractions 0.01-0.9 for the sagital mouse brain dataset.** **A:** From top to bottom, metrics shown are Transcriptomic Hausdorff distance, coordinate Hausdorff distance, ARI, Mean difference between full and sketched PCA matrices, and mean squared error in local neighborhood composition (kNN = 10).

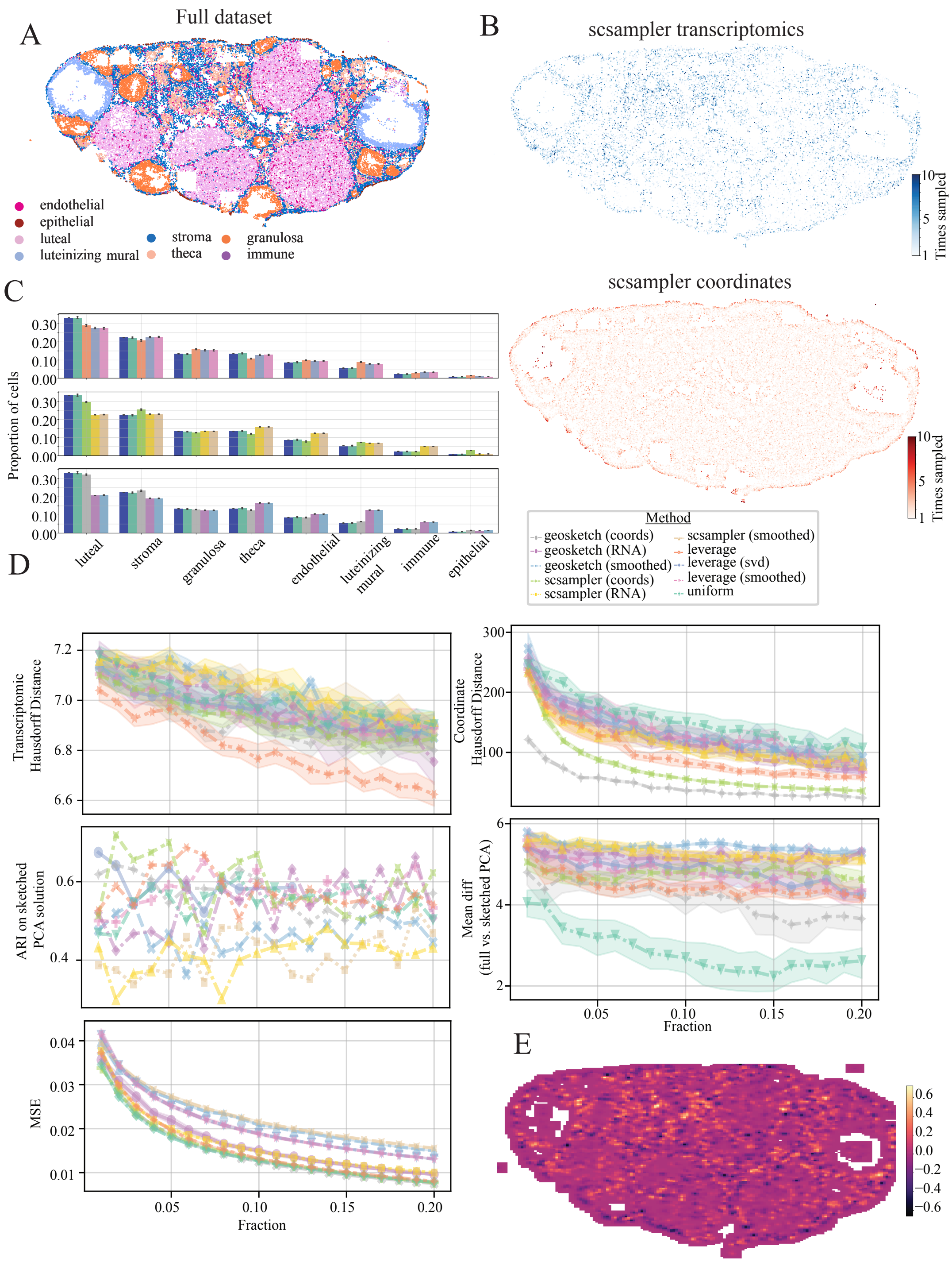

Extended Data Fig. 8: **Mouse Ovary results (Vizgen MERFISH). A:** Ground truth cell type labels for full dataset **B:** Resulting sketched datasets for transcriptomic (top right), coordinate (bottom right) and composite (bottom left) based scSampler sketching. Color scale indicates how many times a cell was sampled out of ten iterations of the sketching algorithm **C:** Cell type proportions for full dataset and 0.1 down-sampled sketches across algorithms evaluated. **D:** Evaluation metrics for sketching algorithms. Transcriptomic Hausdorff distance between sketch and full dataset for sampling fractions from $0.01 - 0.20$ (top left), coordinate Hausdorff distance between sketch and full dataset for sampling fractions from $0.01 - 0.20$ (top right), Adjusted rand index score for sketched cluster solutions compared to ground truth (middle left), mean difference between the full vs sketched data projected onto the principle component loadings (middle right), mean squared error (MSE) between sketched and full dataset's local neighborhood composition. Colors indicate sketching method. **E:** Difference in local neighborhood composition MSE between transcriptomic vs coordinate based scSampler sketching at a 10% sampling fraction, visualized on the tissue. Lighter colors indicate higher transcriptomic MSE compared to coordinate based sketching

A

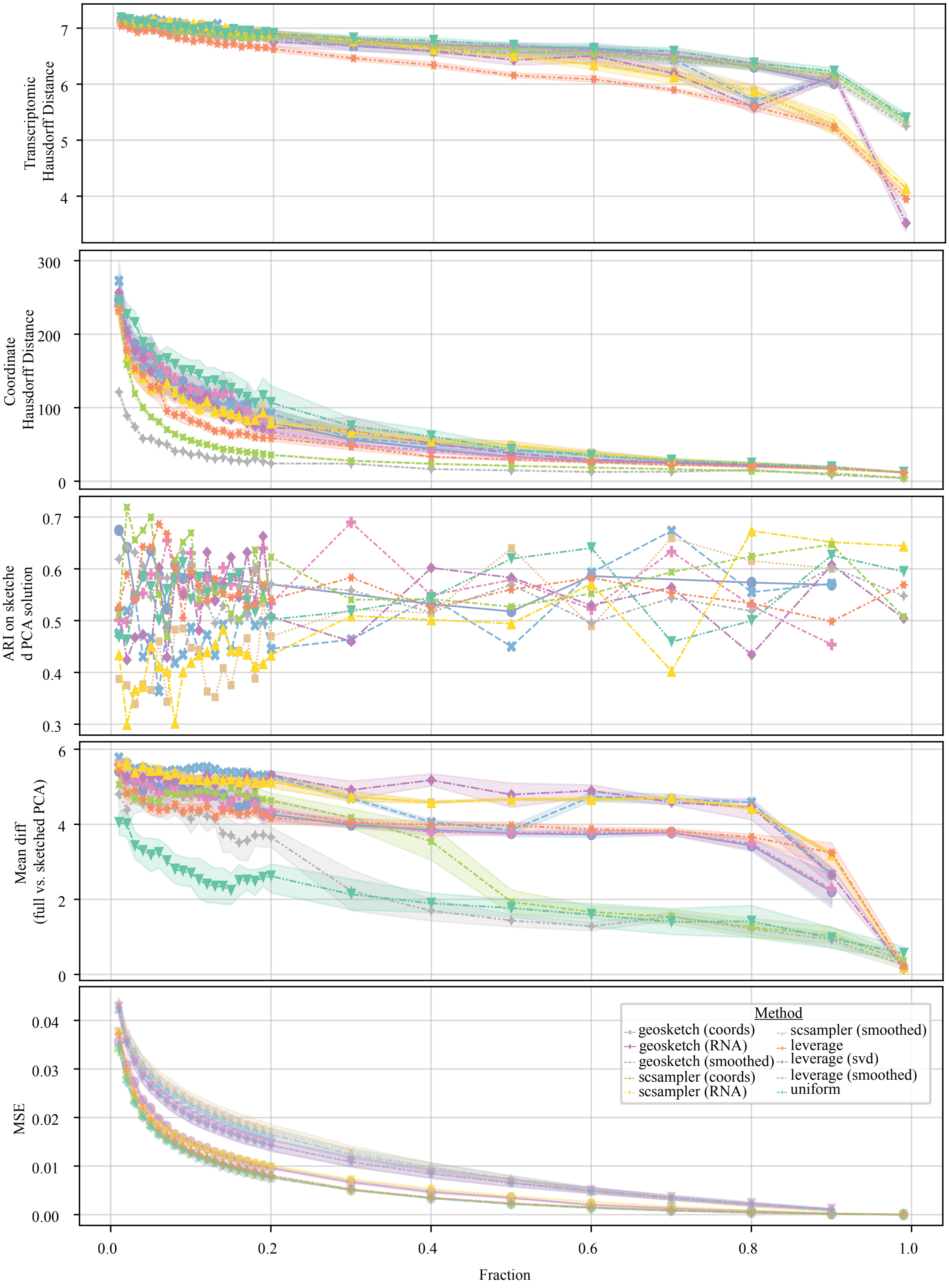

Extended Data Fig. 9: **Sketching evaluation across sample fractions 0.01-0.9 for the mouse ovary dataset. A:** From top to bottom, metrics shown are Transcriptomic Hausdorff distance, coordinate Hausdorff distance, ARI, Mean difference between full and sketched PCA matrices, and mean squared error in local neighborhood composition (kNN = 10).
